# Temporal metacognition as the decoding of self-generated brain dynamics

**DOI:** 10.1101/206086

**Authors:** Tadeusz W. Kononowicz, Clémence Roger, Virginie van Wassenhove

## Abstract

Metacognition, the ability to know about one’s thought process, is self-referential. Here, we combined psychophysics and time-resolved neuroimaging to explore metacognitive inference on the accuracy of a self-generated behavior. Human participants generated a time interval and evaluated the signed magnitude of their temporal production. We show that both self-generation and self-evaluation relied on the power of beta oscillations (β; 15−40 Hz) with increases in early β power predictive of increases in duration. We characterized the dynamics of β power in a low dimensional space (β state-space trajectories) as a function of timing and found that the more distinct trajectories, the more accurate metacognitive inferences were. These results suggest that β states instantiates an internal variable determining the fate of the timing network’s trajectory, possibly as release from inhibition. Altogether, our study describes oscillatory mechanisms for timing, suggesting that temporal metacognition relies on inferential processes of self-generated dynamics.

## INTRODUCTION

Metacognition, the ability to introspect about one’s cognitive state (Fleming & Dolan, 2012), has been little explored in the domain of time perception. Theories of psychological time do not currently account for metacognitive abilities although they are likely fundamental for fine-tuning endogenous timing mechanisms. The capacity to monitor and evaluate one’s action involves some form of time estimation: for instance, we often monitor our timing for appropriate turn taking during conversations. During decision-making, several species can represent timing uncertainties derived from the temporal statistics of external sensory inputs (Mamassian, 2008; Balci et al, 2009; Jazayeri & Shadlen, 2010). Animals can self-monitor their timing (Meck et al., 1984) and humans can reliably judge their timing errors when reproducing a learned target duration (Akdogan & Balci, 2017; also see Miltner et al., 1997). A recent paper extended error monitoring to numerical errors (Duyan & Balci, in press) on the assumption of similar representational features across magnitudes (albeit see Martin et al., 2017). In fact, humans and mice seem to combine their uncertainty estimates of exogenous timing information with endogenous uncertainty regarding how well they performed on a timing task (Balci et al, 2009). Yet, it is unclear whether, in the absence of external timing contingencies provided by sensory stimuli, humans could introspect about timing behavior. In this study, we explored ‘temporal metacognition’ as a self-referential process enabling to render intelligible one’s timing error given a task in which participants self-generated target durations in the absence of prior learning or sensory inputs.

We asked human participants to produce time intervals of 1.45 s, and to subsequently rate their produced interval on a continuous scale going from ‘too short’ to ‘too long’ (**Fig. 1**). In this temporal production task, participants self-initiated the time interval by button press (R1), and terminated it with a second button press when they considered that 1.45 s had elapsed. Participants had full volitional control over their time production in the absence of any sensory cues, which enabled to assess how internal timing become explicitly available to awareness through self-referential metacognition (Block, 1995; Fleming et al., 2012). This experimental design also provided several paradigmatic and conceptual benefits: first, the task requirements capitalized on forward-inverse models in motor execution (Miall & Wolpert, 1998) considering that the key variable for the motor goal was the timing of the second button press given *when* the first button press was realized. This internal variable could be set early during a trial, and used to guide motor trajectory of the second button press allowing the minimization of execution errors (Harris & Wolpert, 1998) *in time*. Second, the metacognitive assessment of the motor timing goal could parsimoniously rely on the availability of such internal variable coding for target duration. Hence, one working hypothesis was that the same internal variable would support temporal production (FOJ: first order judgment) and self-evaluation (SOJ: second order judgment) (Fleming et al., 2012).

**Figure 1.**
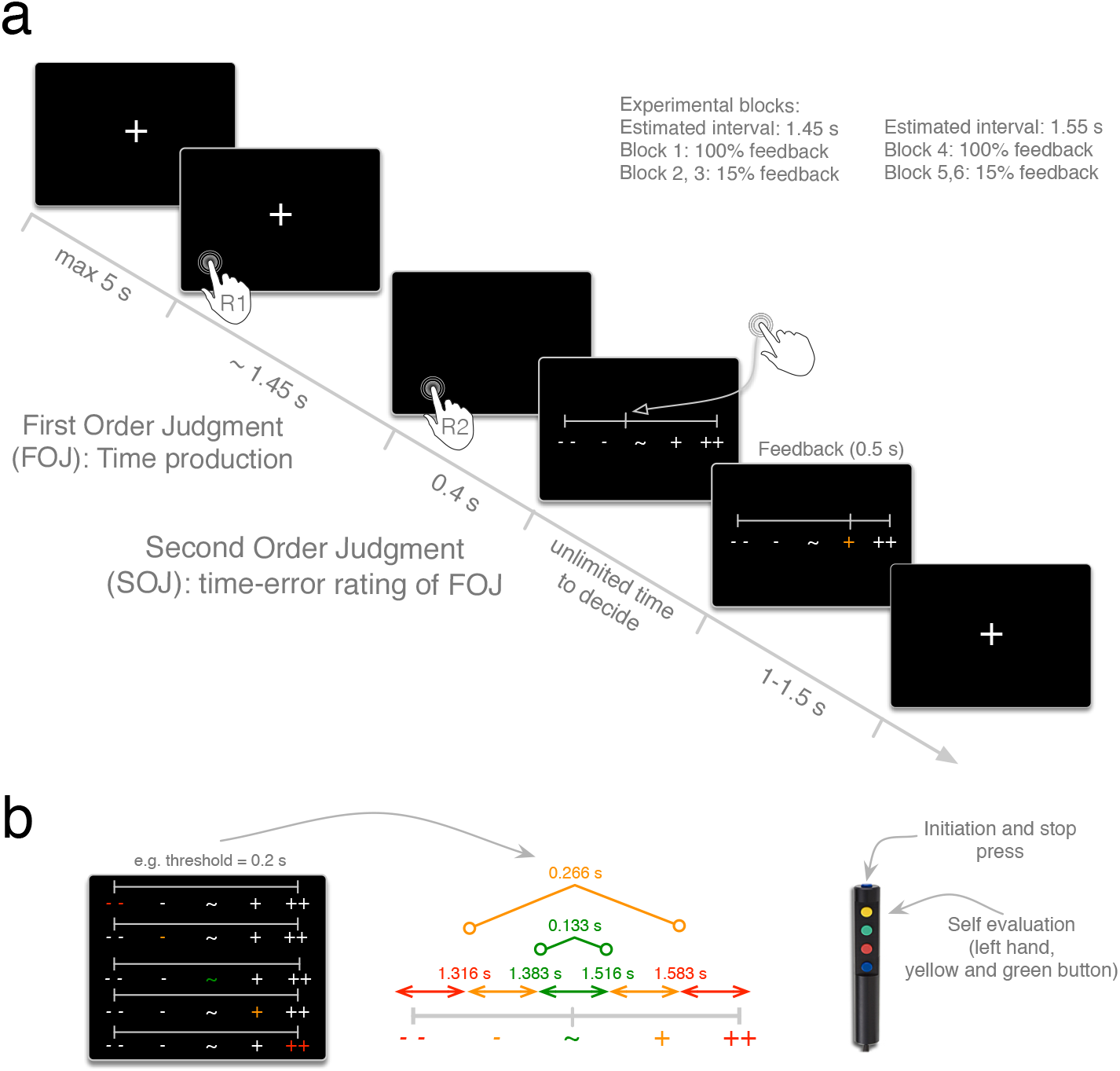
Experimental paradigm in which participants produced a time interval and subsequently estimated its signed magnitude. (**a**) Time course of an experimental trial. (**b**) Rating scale for participants’ self-evaluation and for feedback. Feedback categories were individually tailored according to an individuals’ duration discrimination threshold (green: correct; yellow: slightly shorter or longer; red: too short or too long). The response mapping for TOJ and SOJ is provided on the right.

Current neuroscientific models posit that internal dynamics in timing are mediated by oscillatory or state-dependent network dynamics (Laje & Buonomano, 2013; Buhusi & Meck, 2005; Merchant et al., 2013; Allman et al., 2014; Gu et al., 2015; van Wassenhove, 2016; Bueno et al., 2017). Participants were recorded with combined magneto- and electro-encephalography (MEG, EEG) while performing the task. We quantified the dynamics of oscillatory brain responses when they self-generated (FOJ) and self-evaluated (SOJ) the target duration. In the context of a two-staged timing model (van Wassenhove, 2009), and following a series of studies suggesting that duration may be coded by the power of beta (β) oscillations (Kononowicz & Van Rijn, 2015; Kulashekhar et al., 2016; Wiener et al, 2018), we hypothesized that β oscillations may implement the internal variable coding for duration in the temporal production (FOJ) task. Additionally, we tested whether such internal variable could naturally provide an estimate of the under- or over-estimation of the FOJ for the metacognitive evaluation (SOJ).

We report that participants reliably estimated the direction and the signed magnitude of their timing errors prior to receiving any feedback on the validity of their produced time interval. These results confirmed the ability to detect and estimate one’s temporal errors during self-generated time production. Second, we describe several neural markers tracking participants’ first and second order judgments: the power of β oscillations linearly related to the produced duration (FOJ) and non-linearly to metacognitive inference (SOJ). Crucially, the distance in β state-space informed on the reliability of an individual’s metacognitive inferences. Altogether, our findings provide novel evidence for the neural signatures of estimation of self-generated internal variables in motor timing.

## METHODS

### Participants

Nineteen right-handed volunteers (11 females, mean age: 24 years old) with no self-reported hearing/vision loss or neurological pathology were recruited for the experiment and received monetary compensation for their participation. Prior to the experiment, each participant provided a written informed consent in accordance with the Declaration of Helsinki (2008) and the Ethics Committee on Human Research at Neurospin (Gif-sur-Yvette). The data of seven participants were excluded from the analysis due to the absence of anatomical MRI, technical issues with the head positioning system during MEG acquisition, abnormal artifacts during MEG recordings, and two participants not finishing the experiment. These datasets were excluded *a priori* and were not visualized or inspected. Hence, the final sample comprised twelve participants (7 females, mean age: 24 y.o.). All but two participants performed six experimental blocks; the first block was removed for one participant due to excessive artifacts, the last block was removed for another participant who did not conform to the requirements of the task.

### Stimuli and Procedure

Before the MEG acquisitions, participants were explained they were taking part in a time estimation experiment, and written instructions were provided explaining all steps of the experimental protocol. In a given trial, the participant had to perform three successive steps: first, the participant produced a 1.45 s time interval; second, they self-estimated their time production as too short or too long as compared to the instructed time interval and third, they received feedback on their produced time interval (**Fig. 1a**). We will refer to the produced time interval as the first order temporal judgment (FOJ), and to the self-evaluation of the first order judgment as second order temporal judgment (SOJ). The feedback participants received was for all trials in the 1^st^ and in 4^th^ experimental block, or on 15% of the trials in the other blocks (**Fig. 1a**). To tailor an accurate feedback for each individual, a perceptual threshold for duration discrimination of the same 1.45 s duration was collected before the experiment; this individual threshold was used to scale the spacing of the feedback categories as too short, correct or too long (**Fig. 1b**) as well as change the feedback unbeknownst of the participant in blocks 4 to 6.

Each trial started with the presentation of a fixation cross “+” on the screen indicating participants they could start whenever they decided to (**Fig. 1a**). The inter-trial interval ranged between 1 s and 1.5 s. Participants initiated their production of the time interval with a brief but strong button press once they felt relaxed and ready to start. Once they estimated that a 1.45 s interval had elapsed, they terminated the interval by another brief button press. To initiate and terminate their time production (FOJ) participants were asked to press the top button on Fiber Optic Response Pad (FORP, Science Plus Group, DE) using their right thumb (**Fig. 1b**). The “+” was removed from the screen during the estimation of the time interval to avoid any sensory cue or confounding responses in brain activity related to the FOJ. Following the production of the time interval, participants were asked to self-estimate their time estimation (second order judgment; **Fig 1b**). For this, participants were provided with a scale displayed on the screen 0.4 s after the keypress that terminated the produced time interval. Participants could move a cursor continuously using the yellow and green FORP buttons (**Fig. 1b**). Participants were instructed to place the cursor according to how close they thought their FOJ was with respect to the instructed target interval indicated by the sign ‘~’ placed in the middle of the scale. Participants placed the cursor to indicate whether they considered their produced time interval to be too short (‘--‘, left side of the scale) or too long (‘++’, right side of the scale). Participants were instructed to take as much time as needed to be accurate in their SOJ and there was no time limit imposed on participants.

Following the completion of the SOJ, participants received feedback displayed on a scale identical to the one used for SOJ. The row of five symbols indicated the length of the just produced FOJ (**Fig. 1a)**. The feedback range was set to the value of the perceptual threshold estimated on a per (mean population threshold = 0.223 s, SD = 0.111 s). A near correct FOJ yielded the middle ‘~’ symbol to turn green; a too short or too long FOJ turned the symbols ‘-‘ or ‘+’ orange, respectively (**Fig. 1b**); a FOJ that exceeded these categories turned the symbols ‘- -‘ or ‘++’ red. In Block 1 and 4, feedback was provided in all trials; in Block 2, 3, 5 and 6, feedback was randomly assigned to 15% of the trials (**Fig. 1a**). From Block 4 on, and unbeknownst to participants, the target duration was increased to 1.45 + (*individual threshold*/2; mean population duration = 1.56 s). In Block 1 and 4, participants had to produce 100 trials; in Block 2, 3, 5, and 6, participants produced 118 trials. Between the experimental blocks, participants were reminded to produce the target duration of 1.45 s as accurately as possible and to maximize the number of correct trials in each block.

### Estimation of temporal discrimination threshold

The psychoacoustics toolbox was used to calculate the temporal discrimination threshold for each participant (Soranzo & Grassi, 2014) by adapting the available routine “DurationDiscriminationPureTone” provided in the toolbox. An adaptive procedure was chosen using a staircase method with a two-down one-up rule, and stopped after twelve reversals (Levitt, 1971). For each trial, three identical tones of 1 kHz were presented to the participant. One of the tones lasted longer than 1.45 sec (deviant tone) while the other 2 tones lasted precisely 1.45 sec (standard tones). The position of the deviant tone changed randomly across trials. The task was to identify the deviant tone and to give its position in the sequence. Tones were provided by earphones binaurally. The value of the correct category was set as *target duration* +/– (*threshold*/3), the lower and upper limit values were set as *target duration* +/– (2* *individual threshold*/3). These values were used to provide feedback to participants.

### Simultaneous M/EEG recordings

The experiment was conducted in a dimly-lit, standard magnetically-shielded room located at Neurospin (CEA/DRF) in Gif-sur-Yvette. Participants sat in an armchair with eyes open looking at a screen used to show visual stimuli using a projector located outside of the magnetically shielded room. Participants were asked to respond by pushing a button on a FORP response pad (Science Plus Group, DE) held in their right hand. Electromagnetic brain activity was recorded using the whole-head Elekta Neuromag Vector View 306 MEG system (Neuromag Elekta LTD, Helsinki) equipped with 102 triple-sensors elements (two orthogonal planar gradiometers, and one magnetometer per sensor location) and the 64 native EEG system using Ag-AgCl electrodes (EasyCap, Germany) with impedances below 15 kΩ. Participants were seated in upright position and their head position was measured before each block using four head-position coils placed over the frontal and the mastoid areas. The four head-position coils and three additional fiducial points (nasion, left and right pre-auricular areas) were used during digitization to help with co-registration of the individual’s anatomical MRI. MEG and EEG (M/EEG) recordings were sampled at 1 kHz and band-pass filtered between 0.03 Hz and 330 Hz. The electro-occulograms (EOG, horizontal and vertical eye movements), -cardiograms (ECG), and -myograms (EMG) were recorded simultaneously with MEG. EMG recordings were taken from the *flexor pollicis brevis*, which is involved in the thumb flexion. A pair of surface bipolar electrodes were placed on the surface of right thumb (*thenar eminence*) on right hand of each subjects approximately 5 mm apart. The head position with respect to the MEG sensors was measured using coils attached to the scalp. The locations of the coils and EEG electrodes were digitized with respect to three anatomical landmarks using a 3D digitizer (Polhemus, US/Canada). Stimuli were presented using a PC running Psychtoolbox software (Brainard, 1997) that has been executed in Matlab environment.

## Data Analysis

### M/EEG data preprocessing

Signal Space Separation (SSS) was applied to decrease the impact of external noise on recorded brain signals (Taulu & Simola, 2006). SSS correction, head movement compensation, and bad channel rejection was done using MaxFilter Software (Elekta Neuromag). Trials containing excessive ocular artifacts, movement artifacts, amplifier saturation, or SQUID artifacts were automatically rejected using rejection criterion applied on magnetometers (55e^-12^ T/m) and on EEG channels (250e^-6^ V). Trial rejection was performed using epochs ranging from −0.8 s to 2.5 s following the first press initiating the time production trial. Eye blinks, heart-beats, and muscle artifacts were corrected using Independent Component Analysis (Bell & Sejnowski, 1995) with mne-python. Baseline correction was applied using the mean value ranging from −0.3 s to −0.1 s before the first key press.

Preprocessed M/EEG data were then analyzed using MNE Python 0.14 (Gramfort et al., 2014) and custom written Python code. For time-domain evoked response analysis, a low-pass zero phase lag FIR filter (40 Hz) was applied to raw M/EEG data. For time frequency analyses, raw data were filtered using a double-pass bandpass FIR filter (0.8 – 160 Hz). The high-pass cutoff was added to remove slow trends which could lead to instabilities in time frequency analyses. To reduce the dimensionality, all evoked and time-frequency analyses were performed on virtual sensor data combining magnetometers and gradiometers into single MEG sensor types using ‘*as_type’* method from MNE-Python 0.14 for gradiometers. This procedure largely simplified visualization and statistical analysis without losing information provided by all types of MEG sensors (gradiometers and magnetometers).

EMG data were filtered with FIR filter (20 – 200 Hz) and rectified, following standard EMG assessment. Similarly to ERF procedures, baseline correction was applied using the mean value ranging from −0.3 s to −0.1 s before the first or the second key press. Artifacts were automatically rejected using rejection criterion applied on EMG channel (2e-2 V).

### M/EEG-aMRI coregistration

Anatomical Magnetic Resonance Imaging (aMRI) was used to provide high-resolution structural images of each individual’s brain. The anatomical MRI was recorded using a 3-T Siemens Trio MRI scanner. Parameters of the sequence were: voxel size: 1.0 x 1.0 x 1.1 mm; acquisition time: 466s; repetition time TR = 2300 ms; and echo time TE= 2.98 ms. Volumetric segmentation of participants’ anatomical MRI and cortical surface reconstruction was performed with the FreeSurfer software (http://surfer.nmr.mgh.harvard.edu/). A multi-echo FLASH pulse sequence with two flip angles (5 and 30 degrees) was also acquired (Jovicich et al., 2006; Fischl et al., 2004) to improve co-registration between EEG and aMRI. These procedures were used for group analysis with the MNE suite software (Gramfort et al., 2014). The co-registration of the M/EEG data with the individual’s structural MRI was carried out by realigning the digitized fiducial points with MRI slices. Using mne_analyze within the MNE suite, digitized fiducial points were aligned manually with the multimodal markers on the automatically extracted scalp of the participant. To insure reliable coregistration, an iterative refinement procedure was used to realign all digitized points with the individual’s scalp.

### MEG source reconstruction

Individual forward solutions for all source locations located on the cortical sheet were computed using a 3-layers boundary element model (BEM) constrained by the individual’s aMRI. Cortical surfaces extracted with FreeSurfer were sub-sampled to 10,242 equally spaced sources on each hemisphere (3.1 mm between sources). The noise covariance matrix for each individual was estimated from the baseline activity of all trials and all conditions. The forward solution, the noise covariance and source covariance matrices were used to calculate the dSPM estimates (Dale et al., 2000). The inverse computation was done using a loose orientation constraint (loose = 0.4, depth = 0.8) on the radial component of the signal. Individuals’ current source estimates were registered on the Freesurfer average brain for surface based analysis and visualization.

### ERF/P analysis

The analyses of evoked-related fields (ERF) and potentials (ERP) with MEG and EEG, respectively, focused on the quantification of the amplitude of slow evoked components using non-parametric cluster-based permutation tests which control for multiple comparisons (Maris & Oostenveld, 2007. This analysis combined all sensors and electrodes into the analysis without predefining a particular subset of electrodes or sensors, allowing to keep the set of MEG and EEG data as similar and consistent as possible. We used a period ranging from 0.3 s to 0.1 s before the first press as a baseline. For the ERF/P analysis the data were low-pass filtered using 40 Hz FIR filter.

### Time-frequency analysis

To analyze the oscillatory power in different frequency bands using cluster based permutation, we used DPSS tapers with an adaptive time window of *frequency/2* cycles per frequency in 4 ms steps for frequencies ranging from 3 to 100 Hz, using ‘*tfr_multitaper’* function from MNE-Python. The time bandwidth for frequency smoothing was set to 2. To receive the desired frequency smoothing, the time bandwidth was divided by the time window defined by the number of cycles. For example, for 10 Hz frequency time bandwidth was 2/0.5, resulting in 4 Hz smoothing. We used a time window ranging from 0.3 s to 0.1 s before the first press as baseline. Statistical analyses were performed on theta (3-7 Hz), alpha (8-14 Hz), beta (β: 15-40 Hz), and gamma bands (41-100 Hz) submitted to spatiotemporal cluster permutation tests in the same way as for evoked response analyses. Both time-frequency and power spectral density (PSD) estimates were computed using discrete spheroidal sequences tapers (Slepian, 1978). PSD estimates were computed in 1 Hz steps.

To analyze the oscillatory power in β frequency (15-40 Hz) on single trials, we used Morlet wavelets as implemented in the *‘single_trial_power’* from MNE-Python with 5 cycles per frequency in 4 ms steps. We chose this parameter to capture only the initial β power elicited by the first button press and not the following preparatory brain responses to the second button press. We used a period ranging from 0.3 s to 0.1 s before the first press as baseline.

### Cluster based statistical analysis of MEG and EEG data

Cluster-based analyses identified significant clusters of neighboring electrodes or sensors in the millisecond time scale. To assess the differences between the experimental conditions as defined by behavioral outcomes, we ran cluster-based permutation analysis (Maris & Oostenveld, 2007), as implemented by MNE-Python by drawing 1000 samples for the Monte Carlo approximation and using FieldTrip’s default neighbor templates. The randomization method identified the MEG virtual sensors and the EEG electrodes whose statistics exceeded a critical value. Neighboring sensors exceeding the critical value were considered as belonging to a significant cluster. The cluster level statistic was defined as the sum of values of a given statistical test in a given cluster, and was compared to a null distribution created by randomizing the data between conditions across multiple participants. The p-value was estimated based on the proportion of the randomizations exceeding the observed maximum cluster-level test statistic. Only clusters with corrected p-value < 0.05 are reported. For visualization, we have chosen to plot the MEG sensor or the EEG electrode of the significant cluster, with the highest statistical power.

### Binning procedure of behavioral and neuroimaging data

All cluster-based analyses were performed on three experimentally-driven conditions defined on the basis of either the objective performance in time production (FOJ: short, correct, long) or the subjective self-evaluation (SOJ: short, correct, long) separately for each experimental block. Before the binning procedure the behavioral data were z-scored on peer block basis to keep the trial count even in each category. Additionally, computing these three conditions within a block focused the analysis on local variations of brain activity as a function of objective or subjective performance. Additionally, to overcome limitations of arbitrary binning, and to capitalize on the continuous performance naturally provided by the time production and time self-evaluation tasks, we also used a single trial approach, which allowed the investigation of interactions between the first and second order terms.

### Generalized additive mixed models analysis

To analyze single trial data we used generalized additive mixed models (Wood, 2017; GAMM). Detailed discussions on the GAMM method can be found in elsewhere (Wood, 2017). Here, we briefly introduce the main advantages and overall approach of the method. GAMMs are an extension of the generalized linear regression model in which non-linear terms can be modeled jointly. They are more flexible than simple linear regression models as they do not require that a non-linear function be specified and the specific shape of the non-linear function (i.e. smooth) is determined automatically. Specifically, the non-linearities are modeled by so-called basis functions that consist of several low-level functions (linear, quadratic, etc.). We have chosen GAMMs as they can estimate the relationship between multiple predictors and the dependent variable using a non-linear smooth function. The appropriate degrees of freedom and overfitting concerns are addressed through cross-validation procedures. Importantly, interactions between two nonlinear predictors can be modeled as well. In that case, the fitted function takes a form of a plane consisting of two predictors. Mathematically, this is accomplished by modeling tensor product smooths. Here, we used thin plate regression splines as it seemed most appropriate for large data sets and flexible fitting (Wood, 2003). In all presented analyses, we used a maximum likelihood method for smooth parameter optimization (Wood, 2011). Besides F and p values computed using Wald test (Wood, 2012), the Supplementary tables contain the information on the estimated degrees of freedom (edf). Edf values can be interpreted as how much a given variable is smoothed. Although, higher edf values indicate more complex splines, all tested models showed linear splines (edf = 1), depicted in the plotted model outcomes in associated figures.

GAMM analyses were performed using the *mgcv* R package (Wood, 2009, version 1.8.12). GAMM results were plotted using the *itsadug* R package (Van Rij et al., 2016, version 1.0.1).

### Behavioral data analysis using generalized additive mixed models

The analysis of behavioral data was performed as in the GAMMs framework as fully described below in the *Single-trial analysis of MEG and EEG data using generalized additive mixed models*, unless stated otherwise in the Results section. Each model was fitted with participant as a random factor. For the Block analysis, Block was included as a fixed factor. For the analysis of metacognitive inference, FOJ was entered as a continuous predictor of SOJ. The validity of the smooth term was assessed for FOJ using a relative model comparison performed using Akaike Information Criterion (AIC) and *χ*^2^ test.

### Single-trial analysis of MEG and EEG data using generalized additive mixed models

Although not widely used, GAMMs have been proven useful for modeling EEG data (Tremblay & Newman, 2015). Contrary to some of the previous studies using GAMMs for modeling of multidimensional electrophysiological data, sensors were not included as fixed effects. Rather, we took a more conservative approach and fitted the same model for every sensor separately. The resulting p-values were then corrected for multiple comparisons using false discovery rate (FDR) correction (Genovese et al., 2002). For plotting purposes, we collapsed the data across significant sensors after FDR correction and refitted the model.

The fitted GAMMs contained random effects term for participant and fixed effects that were based on theoretical predictions. Specifically, the full model had the following specification: *uV/Tesla/power* ~ *FOJ* + *SOJ* + *SOJ accuracy* + *FOJ*SOJ* + *FOJ*SOJ accuracy*. Besides the random term for participants, the model contained smooth terms for the first and second order judgments, *SOJ accuracy* between the first and second order judgment, and the interaction term between *FOJ* and *SOJ accuracy*. Notably, FOJ, SOJ and other predictors were entered as continuous variables in GAMM analyses as opposed to post-hoc experimental conditions which suffered limitations from choosing arbitrary split point in the data.

Although GAMMs have built-in regularization procedures (meaning that they are somewhat inherently resistant against multicollinearity), multicollinearity can been assessed using variance inflation factor (VIF; *fmsb* R package, version 0.5.2). Here, VIF were assessed for the final model and consisted in averaging data from multiple sensors collapsed over a particular variable at hand. None of the VIF values exceeded 1.1, indicating that multicollinearity was unlikely to have had a major influence on the reported findings. Note that Rogerson (2001) recommended maximum VIF value of 5 and the author of *fmsb* recommended value of 10.

Before entering empirical variables in the model, we calculated normalized values or z-scores: trials in which a given variable deviated more than 3 z-scores were removed from further analysis. This normalization was computed separately for every MEG sensor and every EEG electrode. For single-trial analyses of β power in FOJ, we focused on the maximum power within the 0.4 s to 0.8 s period following the first button press. This time window overlapped with the selected time window that was used in cluster analyses. The main difference for the width of the time window is that we used one value for the GAMM and hence insured we capture only the β power elicited by the first button press and not spurious preparatory brain responses to the second button press.

### Linear mixed model analysis of β state space and metacognitive inference

As described above, the β state space was estimated on per block and per participant basis. As splitting the data per block and per individual could inflate the degrees of freedom, we used linear mixed-effects models (e.g., Pinheiro and Bates, 2000; Gelman and Hill, 2007) which, by default, accounted for individual, multiple per subject observations in the data. Linear mixed-effects models are regression models that model the data by taking into consideration multiple levels. Subjects and blocks were entered in the model as random effects that were allowed to vary in their intercept. P-values were calculated based on a Type-3 ANOVA with Satterthwaite approximation of degrees of freedom, using *lmerTest* package in R (Kuznetsowa et al., 2017). The mixed-effects models approach was combined with model comparison that allowed evaluating the best fitting model in a systematic manner.

### Demixed Principal Component Analysis (dPCA)

To extract patterns in different frequency bands of averaged brain activity over experimental conditions, we used dPCA analysis (Kobak et al., 2016). Although, dPCA is commonly used in the population analysis of single cell recordings we apply it to collection of sensors, something that is analogical to population of neurons with mixed selectivities. Detailed discussions and proofs of the dPCA method can be found elsewhere (Kobak et al., 2016). In brief, dPCA aims to find a decomposition of the data into latent components that are easily interpretable with experimental conditions, preserving the original data to a maximal extent. The method compresses the data but also demixes dependencies on measured quantity of the task parameters. dPCA is essentially driven by a trade-off between demixing and compression, thus is a mixture of ordinary PCA and linear discriminant analysis (LDA): PCA aims at determining a projection of the data that optimally separates conditions and LDA aims at determining a projection of the data which minimizes the reconstruction error between the projections and the original data. All analyses were performed using a *Python* version of the *dPCA* module (https://github.com/machenslab/dPCA). To prevent overfitting, we used a regularization procedure to find the optimal λ parameter for our dataset, and used a 20-fold cross-validation. We focused dPCA analyses on the two first components (dPCA 1 and dPCA 2) of the β oscillatory activity because they explained on average 73% (SD= 20%) and 26% (SD= 20%) of the overall variance, respectively. Multidimensional distance was quantified as Euclidean distance for both dPCA analysis and for per-sensor analysis.

### Robust regression of MEG and EEG data

Robust regression is an alternative to least squares regression when data have outliers or influential observations. It is also an alternative to Spearman’s correlation when a more complex model has to be built. Model comparisons were performed using robust F test. All analyses were performed using *robust* R package (Wang et al., 2017, version 0.4-18).

## RESULTS

We first provide behavioral evidence showing that participants can accurately perform a self-generated temporal production (FOJ), and access its precision (SOJ). We then quantified oscillatory signatures of FOJ, and explored the idea of a two-staged process serving metacognitive inferences (SOJ). Our analytical approach used statistical modeling of single-trial data, and dimensionality reduction methods for M/EEG data.

### Drifting time production (FOJ) over time with no loss of precision

In the course of the experiment, participants (*n* = 12) accurately produced the 1.45 s target interval (Blocks 1-3) and the adjusted 1.56 s intervals (Blocks 4-6: feedback was implicitly and individually adjusted to a new target value; cf. Methods) as depicted in the density function over FOJ (**Fig. 2a**). As can be seen in **Fig. 2b**, the increase of FOJ was steady and progressive over the course of the entire experiment. Given the serial task structure, the consecutive changes in feedback (for implicit target durations) co-occurred with an unexpected drift of participants’ duration estimation: in the 100% feedback blocks (Blocks 1 and 4), time intervals were accurately produced with a mean of 1.490 s and 1.582 s, respectively (**Fig. 2a**). In the 15% feedback blocks (Blocks 2, 3 and 5, 6), the length of the FOJ significantly increased (**Fig. 2a**; dashed lines). This was tested by adding the feedback factor in a Generalized Additive Mixed Model (Wood, 2017; GAMM), allowing testing the statistical dependencies on single-trial basis. In these 15% feedback blocks, participants’ temporal productions were significantly longer for the 1.45 s target interval (**Fig 2a**; *T*(8.0) = 11.3, *p* < 10^-15^; +90 ms on average) and for the 1.56 s adjusted interval (*T*(7.6) = 10.4, *p* < 10^-15^; + 78 ms). This suggested a natural tendency of participants to lengthen their time estimates over time when receiving less feedback. However, the blocks providing less feedback (Blocks 2, 3 and 5, 6) systematically occurred after the 100% feedback blocks (Blocks 1 and 4, respectively). Hence, although the amount of feedback was the sole factor to be warranted in the models containing the intercept (1.45 s target: ΔAIC = 123, *χ*^2^(1.0) = 6084832, *p* < 10^-15^; adjusted 1.56 s interval: ΔAIC = 104, *χ*^2^(0.9) = 4500629, *p* < 10^-15^), we could not disentangle with full certainty the time-on-task effect, and the specific feedback effects.

**Figure 2.**
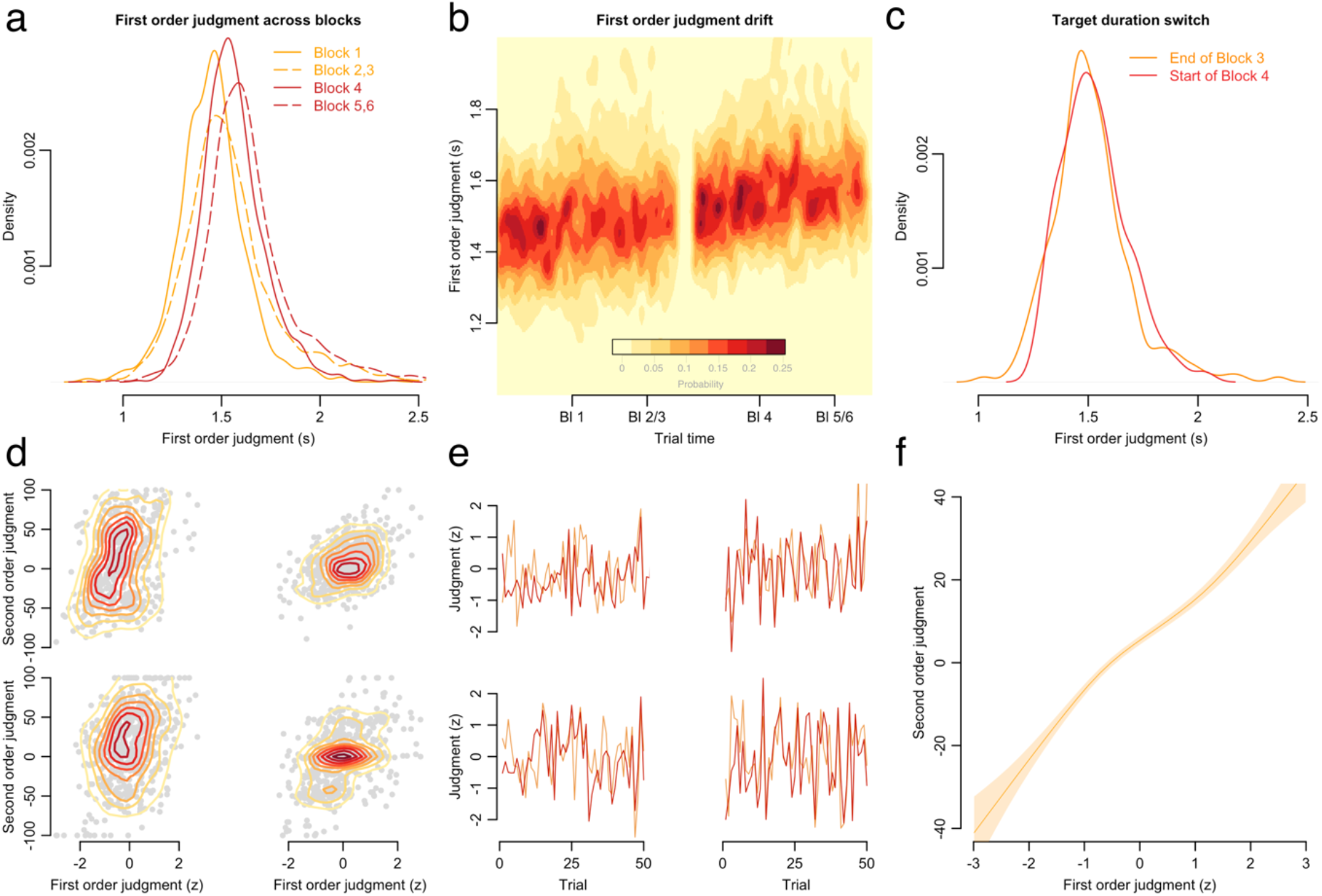
Behavioral evidence for temporal production (FOJ) and participants’ ability to self-evaluate the signed magnitude of produced duration (temporal metacognition, SOJ). (**a**) Probability density function of all participants’ time productions separately for each experimental block. Yellow is 1.45 s; red is 1.55 s. (**b**) Continuous density plot of duration productions in the course of the whole experiment. Red values indicate the distribution of temporal productions for all participants (n=12) throughout all experimental blocks. The yellow break indicates the transition between Blocks 3 and 4. (**c**) Probability density functions of the last 10 trials in Block 3 and the first 10 trials in Block 4, computed across individual trials for all participants. (**d**) Raw data of four participants illustrating SOJ as a function of FOJ in a two dimensional density plot marked by the isometric lines. (**e**) The four participants’ first 50 experimental trials were z-scored for easier visualization: SOJ (orange) tracked FOJ (red) over time. (**f**) The regression line is the GAMM model fit between SOJ and FOJ estimates for all participants.

Nevertheless, additional evidence strongly suggested that the amount of feedback was not the main contributor for the observed drift in time estimation (**Fig 2b**). For instance, to assess the possibility of a step-like transition between Block 3 (explicit 1.45s target) and 4 (implicit adjusted 1.56s target), we used the last 10 trials of Block 3 and the first 10 trials of Block 4. The transition trials from Block 3 and 4 did not significantly differ (**Fig. 2c**; *F*(1) = 0.6, *p* > 0.1) suggesting no clear behavioral transition when the feedback was implicitly changed, even when provided on every single trial (Block 4).

Additionally, given the lengthening of FOJ, we tested whether the precision of time production changed in the course of the experiment. If so, it could have indicated that participants were not performing the task with consistent attentional focus over time, or had changed their cognitive strategy when feedback was implicitly changed. We measured the standard deviation per block and found that, in spite of the drift, the precision of FOJ did not significantly differ across blocks as tested by repeated measures ANOVA (F(5, 65) = 0.7, *p* = 0.634). The stability of behavioral precision throughout the experiment indicated that participants were performing the task as required and that the role of feedback in the task performance was not predominant.

It is thus noteworthy that, despite not explicitly informing participants about the change in target duration introduced in Blocks 4, 5, and 6, participants readily adjusted their temporal production (FOJ) without any loss in the precision of their temporal production (**Fig. 2c**). This observation suggested that humans can implicitly monitor their temporal criterion as previously shown in rats (Meck et al., 1984).

We then asked whether humans could monitor such internal criterion explicitly, as suggested by recent work (Akdogan & Balci, 2017). For this, participants had to introspect about their produced duration (SOJ). Due to the progressive lengthening of FOJ across experimental blocks, the behavioral data were z-scored separately for each block to allow exploiting and analyzing the local variations in single-trial estimates without any confounding influence of time-on-task effects.

### Behavioral evidence for metacognitive inference in time estimation

Individuals could estimate and track their FOJ over time (**Fig. 2d, e**, respectively). To understand how precisely participants could do so, we assessed whether the metacognitive inferences (SOJ) were predictive of self-generated temporal production (FOJ) on a single-trial basis using a Generalized Additive Mixed Model (Wood, 2017; GAMM), which can accommodate nonlinear predictors and interactions. FOJ and SOJ were strongly correlated on a trial-by-trial basis (**Fig. 2f**, *F*(4.0) = 192.5, *edf* = 4.0, *p* < 10^-15^), suggesting that participants could correctly assess the signed error magnitude of the just produced target duration. Additionally, the model fit revealed a nonlinearity in the regression slope between FOJ and SOJ (**Fig. 2f**). This nonlinear term was modeled using a flexible nonlinear spline term provided by GAMM (cf. Methods) and was compared to the model allowing only linear terms (ΔAIC = 15.5, *χ*^2^(4.6) = 18576, *p* < 0.001). This nonlinearity indicated that the correlation was slightly less steep for the FOJ intervals close to the target duration, and steeper for FOJ away from the target duration. Hence, this pattern indicated that FOJ closest to the internal target duration criterion were harder to self-estimate (SOJ). A consistent pattern was also observed in brain activity and will be reported later on.

We then asked whether the SOJ effect resulted from a sequential effect due to the feedback delivered on the previous trial. To test this, we fitted the model adding the factor ‘FOJ on n-1 trial’ as a fixed effect. The analysis was constrained to blocks with 100% feedback (Block 1 and 4) so that all trials - but the first one - were effectively preceded by feedback. The model comparison showed a marginal trend towards an n-1 duration effect (ΔAIC = 0.9, *χ*^2^(1.0) = 2135, *p* = 0.091), suggesting that the longer the previous FOJ was, the shorter the current SOJ tended to be (*F*(1.0) = 2.9, *edf* = 1.0, *p* = 0.09). The trend towards sequential effect in duration production was suggestive of implicit monitoring in humans in line with previous studies (Meck et al., 1984; Meck 1988) showing that rats’ anticipatory behavior was negatively correlated with response times in the previous trial. Nevertheless, on a trial-by-trial basis, the strongest statistical relationship we observed was between FOJ and SOJ (*F*(1.0) = 293.2, *edf* = 1.0, *p* < 10^-15^), suggesting that human participants most exclusively based their self-evaluation (SOJ) on the current trial estimation.

An additional scenario accounting for the interaction between FOJ and SOJ was that participants intentionally increased the variance of their FOJ to improve their SOJ. This cognitive strategy would confound the possibility of an internal state variable being monitored; this possibility could also be an issue in previous work (Akdogan & Balci, 2017). To address this possible confound, we performed a control experiment in which we manipulated the incentives and showed that participants effectively complied with the goal of inferring the signed magnitude of their temporal errors as per task requirements (**Fig. S1a. Control behavioral experiment)**.

To sum up, our behavioral findings indicated that participants produced temporal targets with overall constant precision in the course of the experiment, and that timing errors during temporal production could be assessed through metacognitive inference. We then tested the working hypothesis that the neural markers of the internal variable coding for the target duration may support both first and second order judgments. For this, we analyzed the M/EEG data recorded while participants underwent the behavioral task.

### β power as a state variable for FOJ

Following the initiation of a production trial (R1), we observed a significant cluster of β power in both EEG and MEG activity (*p* = 0.020; p = 0.043; respectively; 0.4-1.2s after R1, **Fig. 3a; Fig. S2ab. Slow brain activity does not encode duration**). When sorting the data according to FOJ, we observed that stronger β power was associated with longer durations (**Fig. 3a)**. To overcome the sorting of brain activity as a function of arbitrary FOJ bins, we also used FOJ as a continuous predictor in a single trial analysis (GAMM). This analysis confirmed that longer durations elicited larger β power in a large cluster of sensors (**Fig. 3b**, **Table S1**) with likeliest neural generators located in bilateral motor and midline cortices (**Fig. 3a**). No other oscillatory responses showed significant changes as a function of temporal production (FOJ, *p* > 0.1).

**Figure 3.**
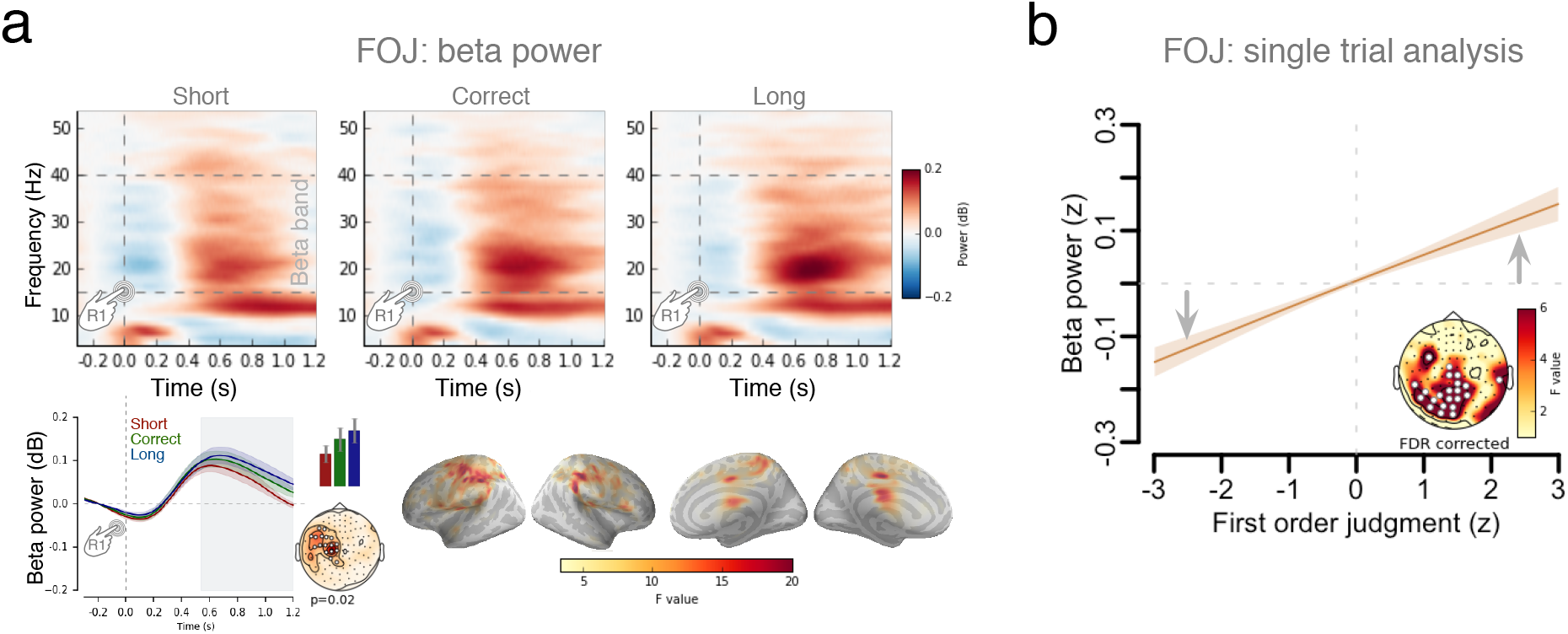
β power as a signature of state variable, controlling produced duration (FOJ). (**a**) Top: time-frequency analysis during FOJ. Left bottom: β (15-40 Hz) power was sorted as a function of FOJ categories (red: too short; green: correct; blue: too long) in the sensor with the highest F-value. The time-based cluster permutation t-test for β power showed latencies (grayed) at which it significantly differed across FOJ categories. The bar plot provides the average β power over significant sensors during that time window. Bars are 2 s.e.m. Right: source estimates contrasting FOJ driven β power effects (uncorrected F-map) implicated medial and motor regions. (**b**) Regression plot showing that β power covaried with FOJ (GAMM model fit). The topographical map shows the distribution of significant sensors. Statistical details are in **Table S1**. The arrows indicate that the main effect was driven by z-scored β values, which significantly differed from their mean value.

Crucially, FOJ did not correlate with EMG activity (**Fig. S3**, *p* > 0.1) or with duration press (measured between the button press and its release) (*p* > 0.1, GAMM), thereby excluding the contribution of low-level factors to the β power effects observed here and which could have confounded the role of β oscillations in time estimation reported in previous work (Kononowicz & Van Rijn, 2015). Nevertheless, we also considered that β power may have been directly related to motor preparation, and not to timing *per se*. Previous research has shown that changes in β power correlated with latency changes caused by the directional uncertainty of motor responses (Tzagarakis, et al., 2010). Here, no changes in direction were necessary as the same button was used to initiate and terminate the target duration, and no other variables than internal timing was required by the task. An alternative hypothesis was that β power changes were a marker of a drift-like accumulation (Simen et al., 2011), so that the peak latency of β power *preceding* the second button press (R2) would differ across produced durations. Such latency differences would predict that, at a given latency near the second button press, β power would significantly differ across produced durations. To test this possible confound, we fitted the GAMM to the mean values of β power locked to the second button press (0.4 s before the second button press). None of the fitted model terms were significant after FDR correction in either MEG or EEG data (**Fig. S4**), providing no substantial evidence for the contribution of motor preparation or accumulation-like processes in the fluctuations of β power.

These results replicated and strengthened the causal implication of β oscillations in duration estimation (Kononowicz & Van Rijn, 2015; Kulashekar et al., 2016; Wiener et al., 2018), here in a motor production task. Covariation of β power with FOJ indicated that this internal timing variable may contribute to the uncertainty of the *“when”* in motor execution. This result is consistent with reports of β power in motor execution tasks controlling for higher level variables (Tan et al., 2016) and the fluctuations of β power as a function of predictability in motor timing (Fujioka et al, 2012; Meijer et al., 2016; Tzagarakis, et al., 2010). Across many studies (e.g., Bartolo et al., 2014; Kononowicz & Van Rijn, 2015; Meijer et al., 2016), β power systematically reached significance ~500 ms following the onset of a timed interval as was also seen here following R1 (**Fig. 3a**); the increase in β power continued thereafter. This pattern suggested the possibility of an early separation of distinct brain states that would be predictive of distinct timing goals inferred during metacognitive evaluation. To explore how β power could be used for metacognitive inference, we assessed the relation between β power, FOJ and SOJ.

### Non-linear use of β power in metacognitive inference

One working hypothesis was that the availability of an internal variable coding for duration may be shared by FOJ and SOJ. We thus asked whether β power varied as a function of SOJ. However, the analysis of β power showed no significant clusters, whether splitting SOJ in categories (**Fig. S5,** all *p* > 0.1) or using SOJ as a continuous predictor in a GAMM (*p* > 0.1). We then reasoned that β power may only be a relevant indicator of SOJ when participants accurately self-estimated their temporal production. To test this, we defined an accuracy score for SOJ as the absolute difference score between FOJ and SOJ. We inverted the scale such that a high SOJ accuracy score indicated that the participant’s metacognitive inference was close to his or her actual temporal production. The full GAMM model allowing non-linear terms had the following specification: β *power* ~ *FOJ* + *SOJ* + *SOJ accuracy* + *FOJ*SOJ* + *FOJ*SOJ accuracy*. FOJ, SOJ, and other predictors were continuous variables (cf. Methods). Although the main term SOJ accuracy and the interaction between FOJ and SOJ showed no significant clusters, the interaction between FOJ and SOJ accuracy revealed a significant change of β *power* in selected sensors (**Fig. 4**, **Table S2**). The nonlinear interaction between FOJ and SOJ accuracy was significant as compared to a model that did not include this interaction term (**Fig. 4**; ΔAIC = 14.0 *χ*^2^(2.1) = 9.6, *p* < 0.001). In sum, the use of GAMM showed that β power was non-linearly related to SOJ accuracy with increased β power for trials in which participants provided an accurate metacognitive inference on their temporal production, *i.e.*, when participants were aware of the direction and of the magnitude of their time errors. The weakened association between β power and temporal production suggested that, on trials with lower SOJ accuracy, the internal variable manifested by β power was more difficult to discriminate. In other words, trials with a distinctive pattern of β power should be more accurately discriminated, an observation consistent with the non-linearities reported in our behavioral results.

**Figure 4.**
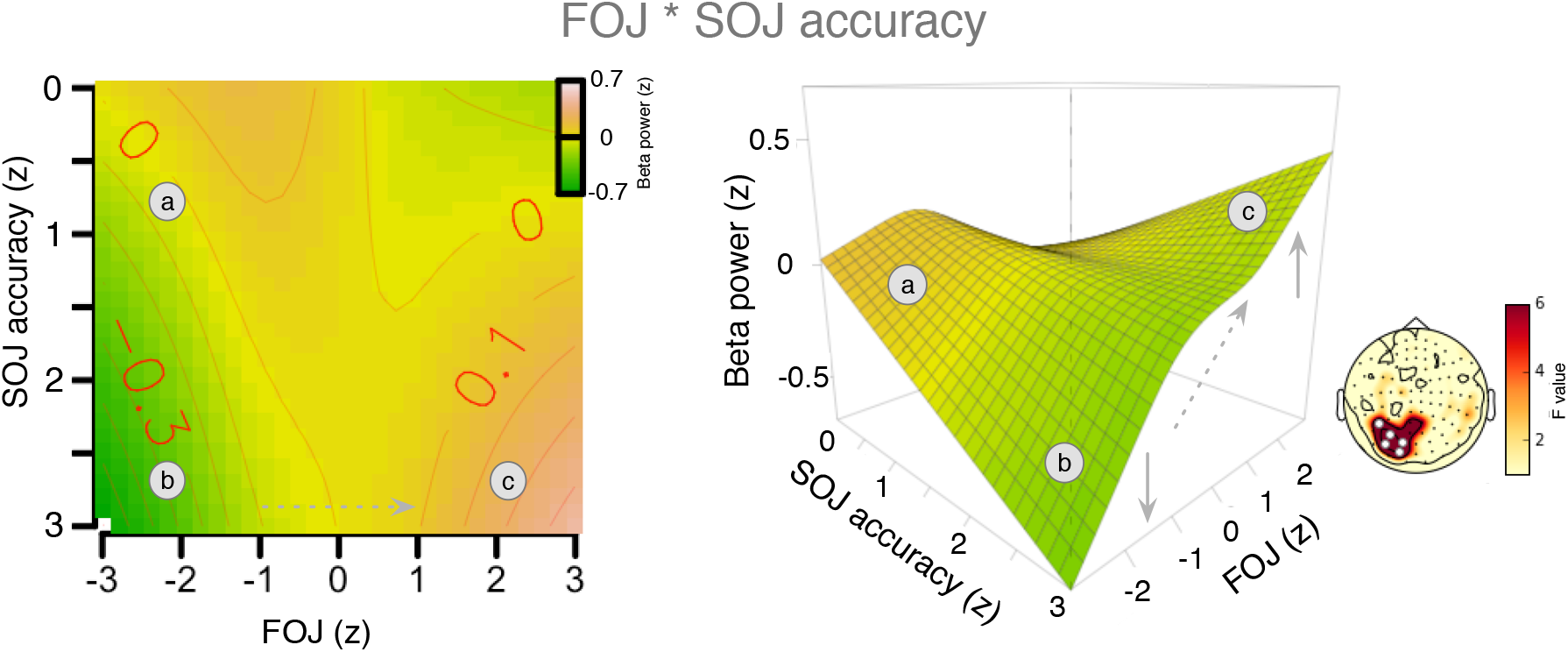
β power is used for metacognitive evaluation. Outcomes of the GAMM model fit for the interaction term between FOJ and SOJ accuracy. The analysis of SOJ accuracy scores indicated that β power was a strong predictor of FOJ but mostly when SOJ were accurate. The 3D plot and the heat map term illustrate that for high SOJ accuracy scores, the power of β oscillations strongly predicted the FOJ. This effect could be seen as an increase of β power from ‘b’ to ‘c’ in the ‘accurate’ part of the surface (**left panel**), indicating that when participants correctly self-estimated their temporal production, β power was also predictive of FOJ. As the SOJ accuracy score decreased towards ‘a’, the predictive power of β for FOJ also decreased. Statistical details in **Table S2**

In light of the relation between FOJ and SOJ accuracy, we thus hypothesized that participants may have access to the internal variable manifested by β power. More distant states of the internal variable would allow more accurate read-out. Therefore, under this working hypothesis, the more distant β power trajectories in the state space, the better metacognitive judgments may be. This was also suggested by our behavioral results (**Fig. 2f**) in which SOJ closest to the FOJ were harder to self-estimate. In light of the non-linearities between FOJ and SOJ and our initial working hypothesis of a common internal variable for FOJ and SOJ, we next aimed to provide a finer assessment of β power. For this, we projected the dynamics of β power into a lower dimensional space using dPCA (Machens, 2010; Kobak et al., 2016) and quantified the distance in this defined ‘β state space’ as a function of FOJ. We specifically asked whether the distinctiveness of β power trajectories was a good predictor of metacognitive inference.

### Distance in β state space predicts metacognitive inference

To project dynamics of β power into lower dimensional state-space with the dPCA method, we used the average power in β band for each individual and each block and across all sensors (from −0.3 to 1.2 s after R1) (Machens, 2010; Kobak et al., 2016). The same approach was used in the *per* individual analyses (**Fig. S6**). Although the dPCA method was designed for single neuron populations, we used it here across sensors, with sensors homologous to a population of neurons with mixed selectivity. We looked at the morphology of the two first demixed Principal Components (dPC1 and dPC2; cf. Methods) which showed that β power rose fast at the onset of the FOJ, and remained segregated in different locations of the state-space (**Fig. 5a**). This pattern is illustrated by plotting the first two dPCs for two example blocks differing in the extent of their state-space separation (**Fig. 5a,** and example subjects **Fig. S6a)**. The early separation pattern was found for nearly all participants in the study (**Fig. S6b**) with an apparent inter-individual variability in the magnitude of the separation in β power state-space (**Fig. S6c)**.

**Figure 5.**
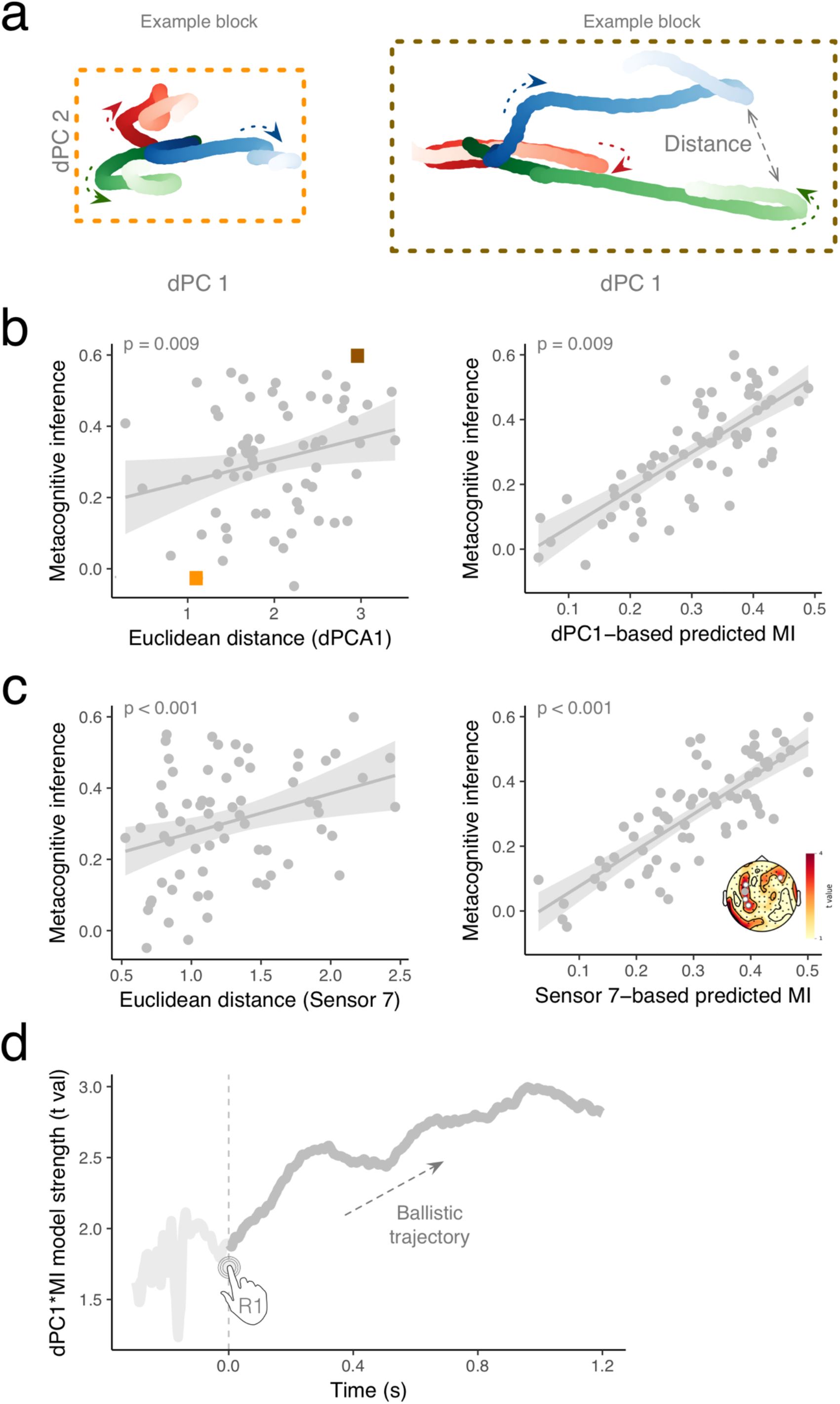
Distance in β state space supports metacognitive inference. (**a**) dPCA decomposition in β band (−0.3 to 1.2 with respect to R1) illustrated for two blocks with low and high metacognitive performance, respectively. The two first dPCs are displayed against each other, and illustrate an early separation of path trajectory. Arrows indicate the progression of β state along the arrow of time. (**b**) The left panel illustrates the significant results of a single trial like analysis in which a linear mixed model was used to predict the level of metacognitive inference (FOJ*SOJ) on the basis of a distance in β state space (dPC1)) on within participant basis. One dot is a block within participant. Orange and grey rectangles correspond to trajectories reported in (a) for two blocks with a low and a high metacognitive inference ability, respectively. The right panel illustrates empirical values of metacognitive inference plotted against the values predicted by the model. (**c**) The left panel illustrates the significant outcome for the same analysis performed on individual sensor (here, one dot is one block, one sensor). The right panel illustrates empirical values of metacognitive inference plotted against the values predicted by the model. The topographical map displays t values together with significant sensors marked as white. The grey sensor is the one displayed in the scatterplot. (**d**) Time course of the linear model predicting metacognitive inference on the basis of distance in β state space (dPC1). The distance in state space is predictive of metacognitive inference early in the onset of the production interval and continues to increase throughout the interval.

To investigate the relationship between distance in β state-space and metacognitive inference, we then employed two approaches: a single trial-like analysis and an analysis over participants. We utilized the mean distance computed for each triplets of samples (too short, correct and too long).

We performed a *per* participant and *per* block analysis using linear mixed models, with subject and block as random effects (*cf.* Methods). We tested whether dPC1 predicted metacognitive inference and found that the degree of β state space separation significantly predicted metacognitive inferences [t(65) = 2.7, p = 0.009] (**Fig. 5b**), indicating that the bigger the distance between the FOJ, the better the metacognitive inference was. For instance, the sample block framed in dark orange (**Fig. 5a,** *right panel*) showed a much larger spread of their β state-space (**Fig. 5b**) along with higher metacognitive inference accuracies (**Fig. 5b)**. Conversely, the individual framed in light orange (**Fig. 5a,** *left panel*) showed a much smaller spread of their β state-space (**Fig. 5b**) and lower metacognitive inference accuracies (**Fig. 5c)**.

The left and right panels plot the empirical values of metacognitive inference against the empirical dPC1 values and the values predicted by the model on the basis of dPC1, respectively. The same analysis quantified on each sensor confirmed this pattern (**Fig. 5c**).

Importantly, we addressed the possibility that alternations of performance precision, which could drive β power, did not rule out the impact of β state-space in metacognitive inference. Thus we insured that the fluctuations in precision (CV) could not account for these observations (**Fig. S7. β state-space was not driven by Coefficient of Variation (Analysis per participant)**). In line with the control behavioral experiment, this analysis confirmed that alternations of performance precision did not rule out the role of β state-space in metacognitive inference.

To test the reliability of these observations, we performed a second similar analysis *per* participant. We quantified metacognitive inference as the correlation (Spearman’s *rho*) between FOJ and SOJ on a per individual basis. We found a significant correlation between distance in β state-space (FOJ: ‘short’, ‘correct’, ‘long’) and the individual’s metacognitive inference (*rho* = 0.75, *p* = 0.005; **Fig. S6c** *left panel*). We insured that (i) the multidimensional distance in β state-space was not driven by the spread of individual FOJ responses as quantified by their Coefficient of Variation (CV, **Fig. S8a, β state-space was not driven by Coefficient of Variation (Analysis per participant)**), and (ii) that no other frequency bands or components were significantly related to metacognitive inference (**Fig. S8 b-c**). To strengthen the outcomes of the dPCA analysis, we also quantified the Euclidean distance in β power separately, for each sensor. With this method, the estimated distance in β space for a cluster of sensors also significantly correlated with metacognitive inference (**Fig. S6c,** *right panel*).

Lastly, to elucidate the dynamics of the β state-space distance and metacognitive inference, we fitted the mixed model predicting metacognitive inference on the basis of dPC1 for each time sample and observed that the predictability of the model monotonically increased over time (**Fig. 5d**). Notably, the build-up already started at the initiation of the interval and evolved as a stable trajectory over time, indicating the separation between the β state space trajectories and their contribution to metacognitive inference.

To sum up, we showed that the larger the distance in β state space, the more accurate the metacognitive inference within participants, and on *a per* individual basis. In other words, trials with a more distinctive pattern of β activity were associated with more accurate metacognitive judgments. This naturally suggests that the distinctiveness of β pattern supports its read-out. The β state-space analysis provided strong evidence that the distance in β state-space could support temporal metacognition by providing a second order estimation of FOJ. The distance in β state-space may reflect a capacity for metacognitive inference of temporal estimation, akin to retrospectively monitoring and reading-out the state of the implicated β network.

## DISCUSSION

Using a newly designed temporal metacognition task with time-resolved neuroimaging and statistical modeling, we investigated how the human brain monitors its self-generated timing. We report several main findings: first, humans can maintain their precision of temporal production over time despite implicit changes and drifts in mean duration values (FOJ). Second, human participants can accurately self-evaluate the signed error magnitude of their temporal productions (SOJ). Third, the power of β oscillatory activity following the initiation of the time interval was an accurate predictor of FOJ, even more so when participants were aware of their time-errors (SOJ). Intriguingly, the distance or distinctiveness in β state-space separating the self-generated time intervals was indicative of the accuracy with which timing performance could be inferred. Altogether, we interpret our findings as supporting the availability of an internal state variable coding for duration, which sets up the goal for a state-dependent trajectory in time production. Our results support the view that metacognitive inferences would consist in the reading out – or the decoding – of internal state variables, consistent with a recent proposal (Fleming & Daw, 2017). In the context of our task, this may possibly rely on forward-inverse models of state-dependent computations in motor systems (Harris & Wolpert, 1998). Below, we review and discuss the main evidence supporting this viewpoint along with the limitations of the current study.

### β power as a marker of state variable coding for duration

We hypothesized an internal state variable informing *when* the second button press should be made given the first button press was made. The post-movement rebound of β oscillations has seminally been proposed to reflect the idling state of motor cortices (Pfurtscheller et al., 1996), possibly sensitive to sensory afferents (Casimi et al., 2001). The observation that the power of β oscillations post-movement was stronger (smaller) for longer (shorter) timed intervals was consistent with the notion that the strength of network inhibition or idling would be predictive of the produced time interval (FOJ). In another words, the stronger the inhibition, the longer the time delay before the next button press.

However, consistent with recent studies indicating that active cognitive components are encoded in the β rebound (Tan et al., 2016), we discuss why β power is not trivially relating to a passive rebound. *First*, we found no significant impact of simultaneous EMG, providing no evidence for the implication of the strength or afferent feedback confounded with time production. *Second*, the variability in β power did not reflect random variance in time production (or explicit variance induced by the participant, cf. control experiment), but the actual time interval that participants were subsequently aware of. In fact, our results suggested that β power was even more telling of an individual’s FOJ when participants correctly self-estimated their FOJ (SOJ). Considering that β power reflects the amount of inhibition in a network (Whittington et al., 2000; Wang, 2010), β power *de facto* provides an estimation of the time delay before the network converges to a desired state, which, in our context, corresponds to the next movement - consistent with general motor inhibition schemes (Duque et al., 2017). *Additionally*, that the power of β oscillations determines the duration of network inhibition (**Fig. 6**) is consistent with the implication of β oscillations in predictive timing (Arnal & Giraud, 2012) and with the notion of an idle state until network updating or reorganization (Engel & Fries, 2010; Spitzer & Haegens, 2017). The power of β has been repeatedly shown to reflect explicit time estimation and prediction (Kononowicz & Van Rijn, 2015; Fujioka et al., 2010; Wiener et al, 2018) not only in motor timing (Bartolo et al., 2015) but also in perceptual duration estimations (Kulashekhar et al., 2015; Spitzer et al., 2014). Hence, we suggest that, because the power of β oscillations is indicative of the strength of inhibition in the network, it also naturally provides a state-dependent variable, which could be used as a duration code.

**Figure 6.**
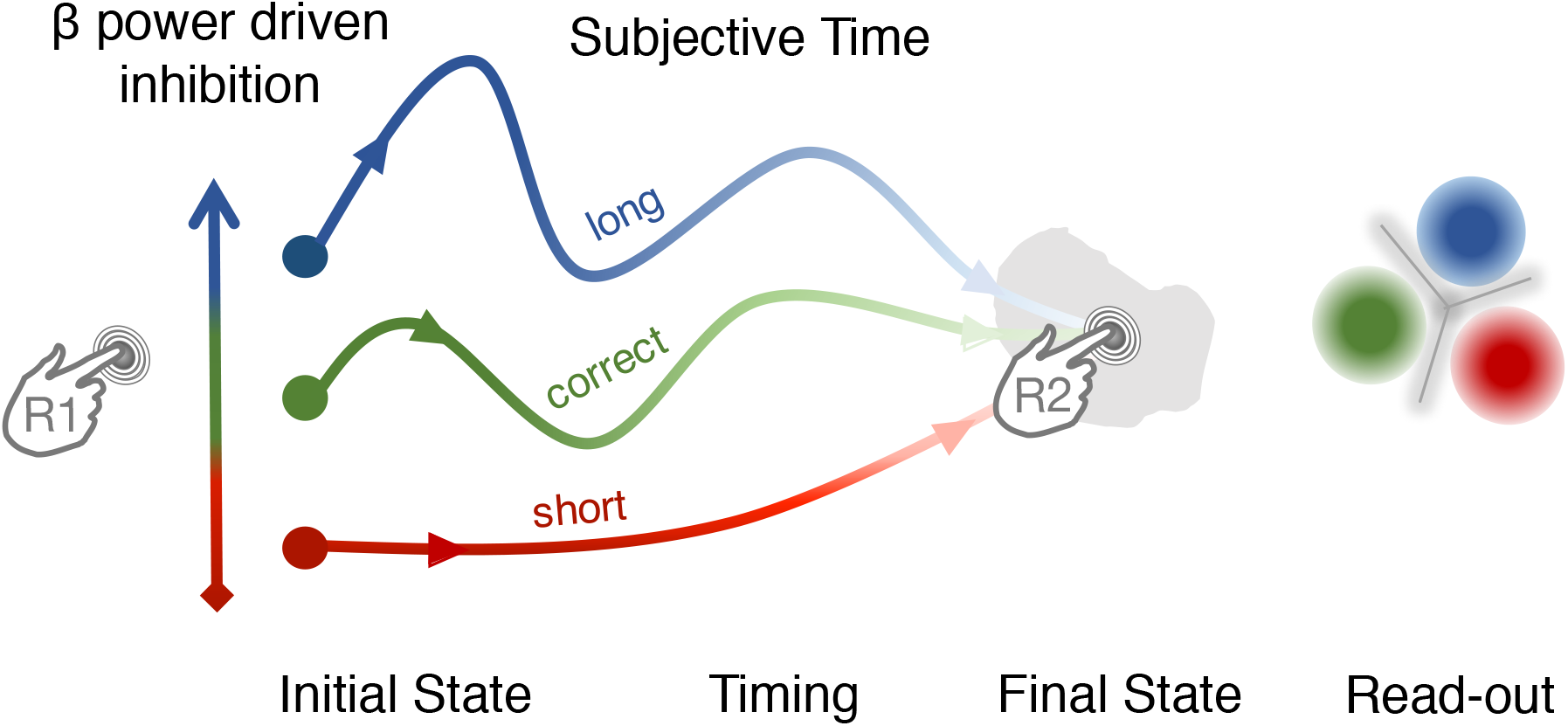
Synthetic view of the proposed role of β oscillations in timing and temporal metacognition. The β power at the onset of the interval determines a network trajectory set by the initial level of inhibition in the network (increasing β power as network inhibition from red to blue). The initial network state and subsequent trajectory determines the motor timing goal. The separation of network trajectories allows reading out the network state associated with a certain motor timing goal.

### Network inhibition and distance in β state space support metacognitive inference

Here, we expand on the notion of inhibition and show how it accounts for two crucial results on temporal metacognition. First, β power was non-linearly related to SOJ with increased β power for trials in which participants provided an accurate metacognitive inference. Second, in the state-space analysis, the more distinct the β state-space trajectories of FOJ categories were, the better participants were at self-evaluating their time errors. The notion of the strength of inhibition departs away from the notion that time estimation relies on the accumulation of sensory or internal evidence (Hardy & Buonomano, 2016; Kononowicz & Penney, 2016; Kononowicz & van Rijn, 2015; van Wassenhove, 2016). The state-variable hypothesis rather suggests that, by indicating the amount of inhibition in the network, β power inherently determines the trajectory that the network will subsequently follow (**Fig. 6**). Hence, the initial state variable associated with the amount of β power may be sufficient to predict the over- or under-duration production, and this was essentially captured by the positive covariation between FOJ and β power in our study. The state variable would be predictive of the cascade of neural events leading to the next volitional button press (**Fig. 6**). What we consider to be the state variable could be formalized as initial inputs into a network, or initial network conditions in the case of a time production task, whose magnitude could control the speed (hence, timing) of the evolution of the system, very much in line with a recent proposal (Wang et al., 2017). By analogy to the β power results, the initial input specifies the position of the initial and final states of the system. Future studies should explore the link between network speed control and β oscillations at the global scale. With respect to the behavioral outcomes, the second press would result from the initial trajectory with limited revisions - or vetoing - shortly before the course of the next action (Schultze-Kraft et al., 2016). Until reaching the state in which the motor plan for the second button press is initialized, the distance in β space would thus reflect the discriminability of state-dependent variable encoding duration and availability for read out, that a decoder or a reader may estimate. That is, the more ‘too short’ trajectory moves away from ‘too long’ trajectory, the likely it is that the resulting duration will be accurately classified. Essentially, we propose that a reader can access the state variable, measured by β power, which determines the fate (trajectory) of network state.

### Conclusions

We showed that the dynamics of β oscillations in the human brain not only predict the self-generation of time intervals but also the metacognitive inferences on their accuracy. Specifically, the distinctiveness of β power trajectories during timing was indicative of metacognitive inference. Our results suggest that network inhibition (β power) instantiates a state variable determining the fate of network trajectory, thus providing a natural code for duration. However, the question remains why one does not correct their ongoing interval production to reduce errors. Future studies will investigate which properties of internally generated representations could be accessible for conscious read out in order to address which mechanisms may support the reading out and the interpretation of the contingencies in the first order network (Cleeremans et al., 2007; Denève et al., 1999; Dayan & Abbott, 2001; Pouget et al., 1998), and whether these mechanisms would be similar to those allowing to read out neural signals that encode primary sensory variables (Komura et al., 2013).

## SUPPLEMENTAL FIGURES

**Figure S1.**
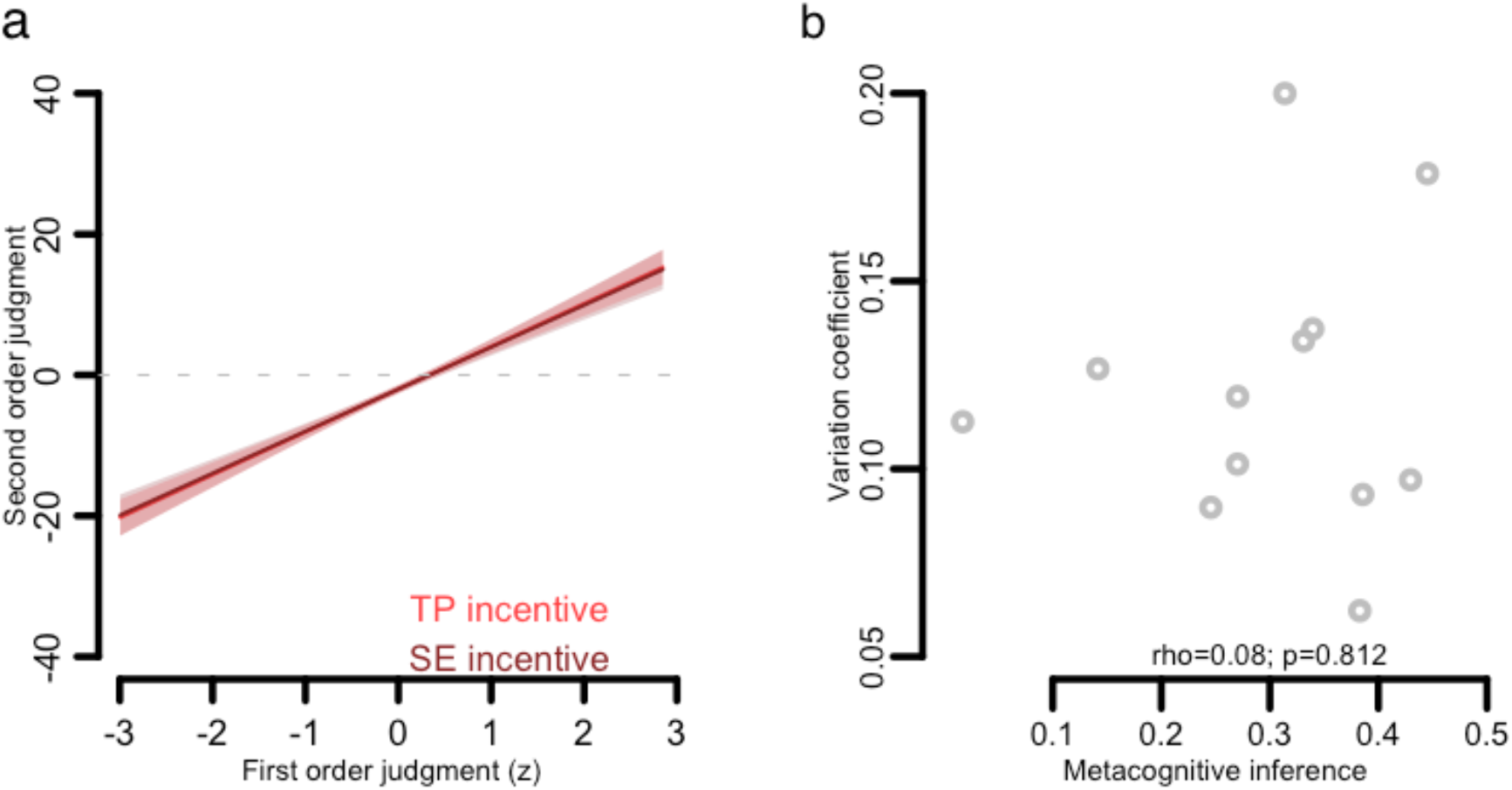
Participants solely relied on temporal representation for their SOJ. (**a**) An additional scenario accounting for the interaction between FOJ and SOJ was that participants intentionally increased the variance of their FOJ to improve their SOJ. This cognitive strategy would confound the possibility of an internal state variable being monitored, and this possibility was also not controlled for in previous work (Akdogan & Balci, 2017). To address this important issue, we performed a control experiment in which we manipulated the incentives: participants produced 1.45 s durations, and received points for their FOJ accuracy in one block or for the congruence between their FOJ and their SOJ in another block. If participants used the confounding cognitive strategy described above, the regression slope for FOJ and SOJ would have been significantly larger in the congruence incentive condition than in the FOJ accuracy incentive condition. In this control experiment, the FOJ significantly covaried with SOJ (*F*(1.0) = 91.9, *edf* = 1.0, *p* < 10^-15^) just as we observed in our main experimental design. Most importantly, the addition of the block type factor was not justified in the model (ΔAIC = 0.9, *χ*^2^(1) = 0.6, *p* = 0.79), demonstrating that participants performed the task sequentially by first estimating FOJ and then estimating SOJ without alternative cognitive strategies. To further strengthen the observations in the control experiment, we checked whether the spread of FOJ estimations could account for an individual’s metacognitive inference. Individual differences in metacognitive inference (FOJ*SOJ) were not correlated with the individuals’ coefficient of variation observed in FOJ (*rho* = 0.08; *p* = 0.81). This observation did not support the idea that participants boost their FOJ and SOJ correlation, but rather that participants effectively complied with the original task goal of inferring the signed magnitude of their temporal errors following their temporal productions. Participants did not modulate their responses based on changes of incentives. The plot depicts behavioral data in the control psychophysical experiment and illustrates the strong link between FOJ and SOJ. Regression lines depict the model fit between the observed SOJ and FOJ estimates for a block with incentive for FOJ (dark red) and another block with an incentive on congruence between FOJ and SOJ (light red). We found no evidence supporting the hypothesis that incentive manipulation impacted SOJ, suggesting that participants in the main task did not use any strategies when performing SOJ. In the incentive for FOJ block, participants received 2 points for a “green” feedback and 1 point for an “orange” feedback. In the incentive for SOJ accuracy block, the distance between FOJ and SOJ category was computed. Participants were given points on each trial according to this distance: the SOJ category boundaries were arbitrary set to −60, −30, +30, and +60 points on the self-evaluation scale. Specifically, when FOJ and SOJ landed in the same category, participants received 2 points; they received 1 point when they landed in adjacent categories; and 0 points when categories were different. Ten participants took part in the experiment with two blocks of 100 trials. The perceptual threshold for duration production on the basis of which the feedback ranges were established in the main experiment, was set to 0.2 for all participants. Participants received feedback in 100% of trials. All other aspects of the control experiment were identical to the main experiment. (**b**) The spread of FOJ was not related to metacognitive inference. The plot illustrates coefficient of variation over FOJ plotted against metacognitive inference quantified as Spearman’s correlation between FOJ and SOJ.

**Figure S2.**
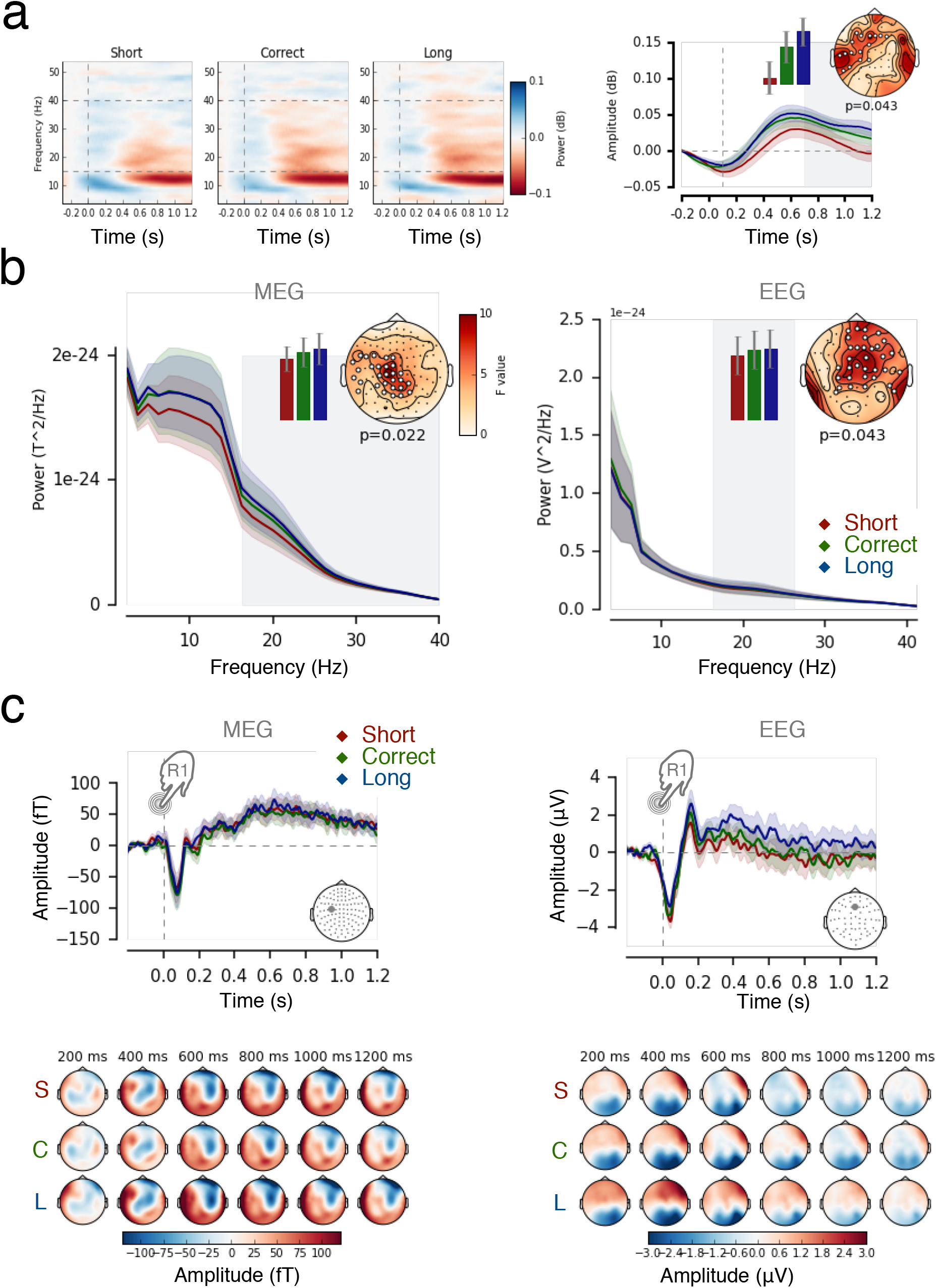
β power, not slow brain activity, controls FOJ. (**a**) We explored the association of slow activity (Contingent Negative Variation, CNV (Kononowicz & Penney, 2016; Macar & Vidal, 2004) with ‘too short’, ‘correct’ and ‘too long’ categories of FOJ. Using data between 0.4 and 1.2 s following the first button press, we found no significant differences in the amplitude of the slow evoked responses as a function of duration estimation (all *p* > 0.1). This replicated previous work discussing the functional implication of slow activity in timing (Kononowicz & Van Rijn, 2011; 2014). Evoked related fields/potentials (ERF/P) following the first keypress showed no significant effects as a function of FOJ in MEG or EEG (left and right panel, respectively). (**b**) The same time-frequency analysis in the same frequencies as those used for MEG analysis (15-40 Hz) confirmed the main MEG observations, showing that larger values of β power yield longer duration productions. (**c**) Power Spectrum Density (PSD) of MEG activity as a function of FOJ. Cluster-based permutation F-test on PSD estimates in the frequency dimension showed significant β (15-40 Hz) power increases as a function of longer duration productions in both MEG and EEG. The vertical grey area demarcates the significant frequency range. The topographical plot illustrates the distribution of sensors included in the cluster. The barplot shows the averaged value in the significant region for all three FOJ conditions in the most significant sensor. EEG analysis, depicted in the right panel, confirmed observations obtained with MEG PSD spectrum for the example electrode.

**Figure S3.**
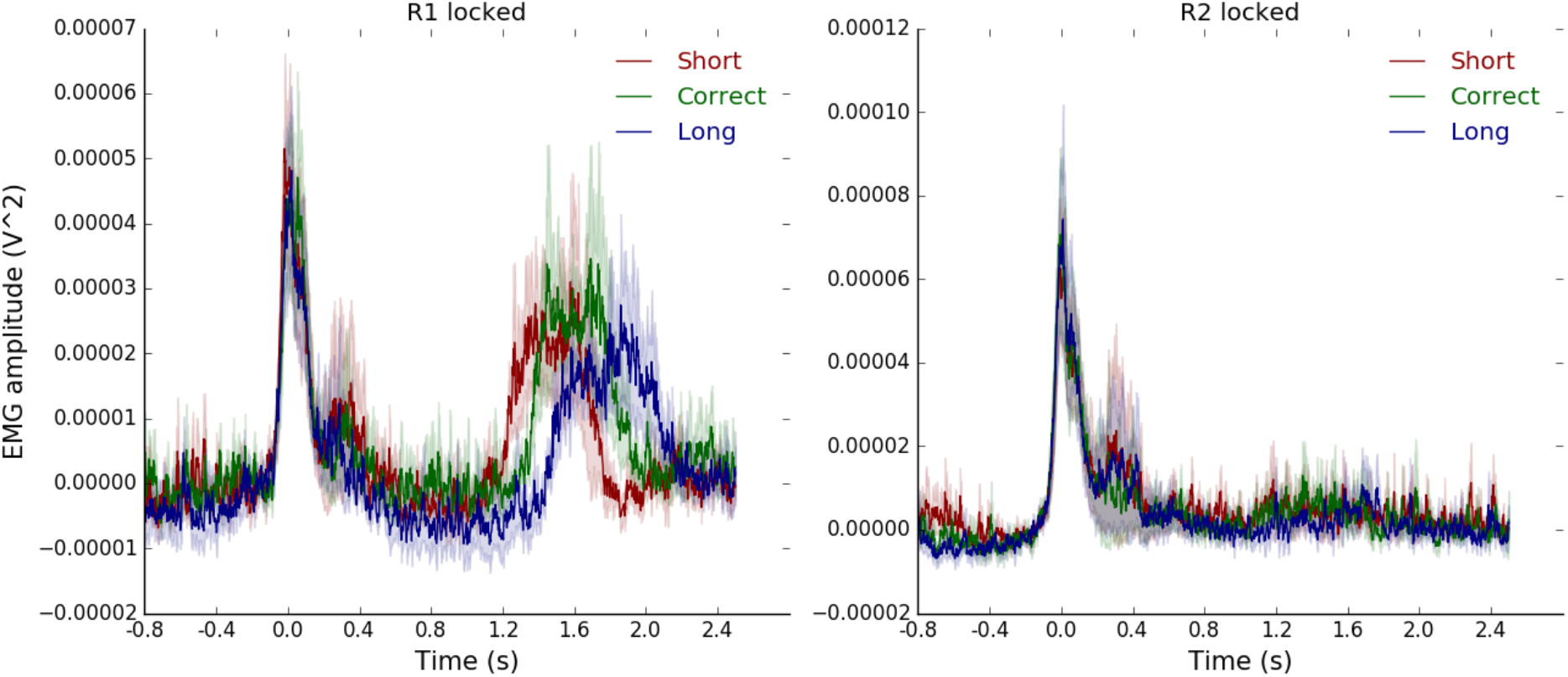
Muscular activity did not impact FOJ. Both panels depict electromyographic (EMG) results. EMG data were split as a function of FOJ performance and time-locked to the first keypress. Differences across time production performances were tested using a one-way non-parametric Friedman’s ANOVA in the 0 to 0.8 s time window following the first keypress with non-overlapping time windows of 0.1 s. We found no significant differences (R1-locked, all *p* > 0.1). The same analysis following the second keypress showed no significant differences between conditions (R2-locked, all *p* > 0.1). Overall, we found no evidence suggesting that motor activity could be a confounding factor for the main M/EEG findings.

**Figure S4.**
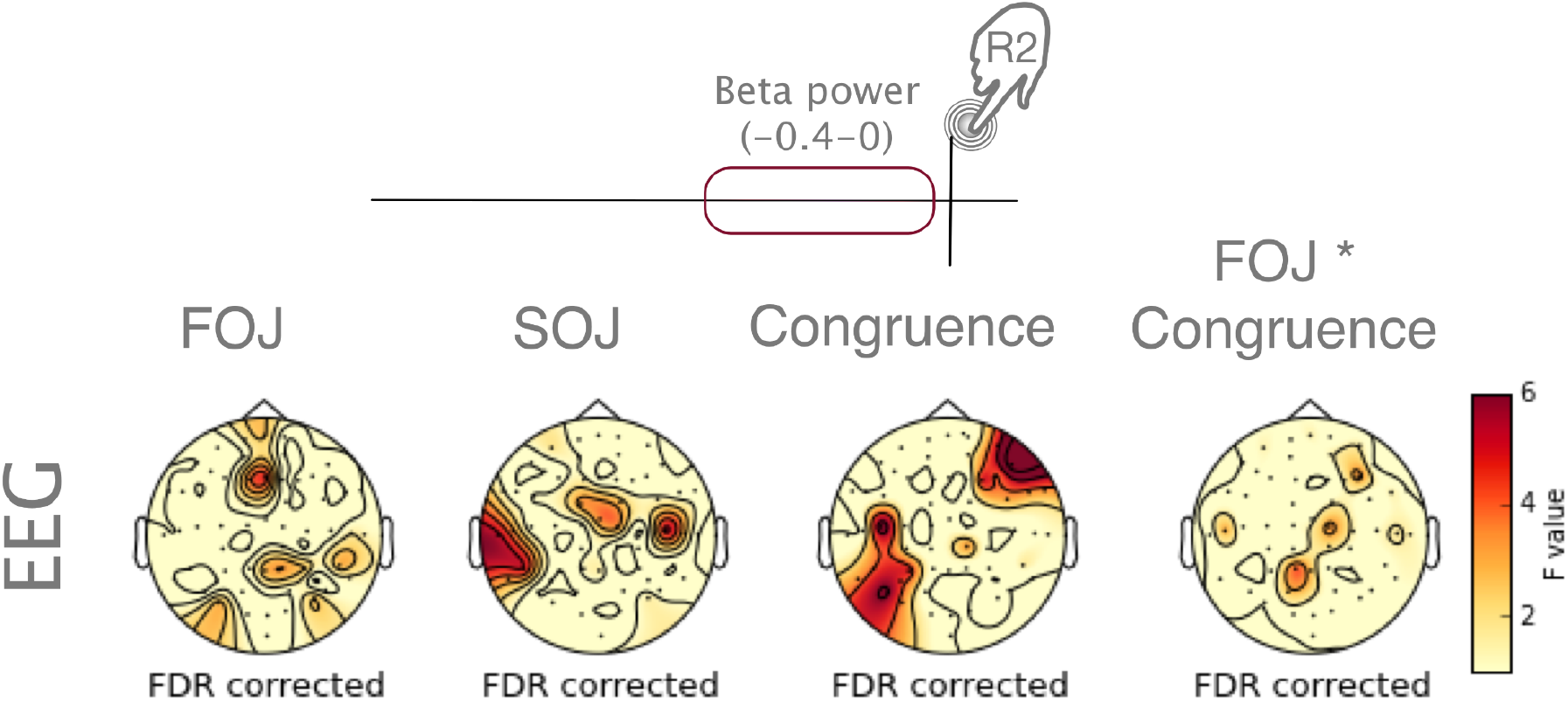
GAMM model fits computed for β power in the 0.4 s time window preceding the second keypress. The topographical maps show that no significant effects were found, suggesting that the second key press was executed after β power reached a certain fixed level.

**Figure S5.**
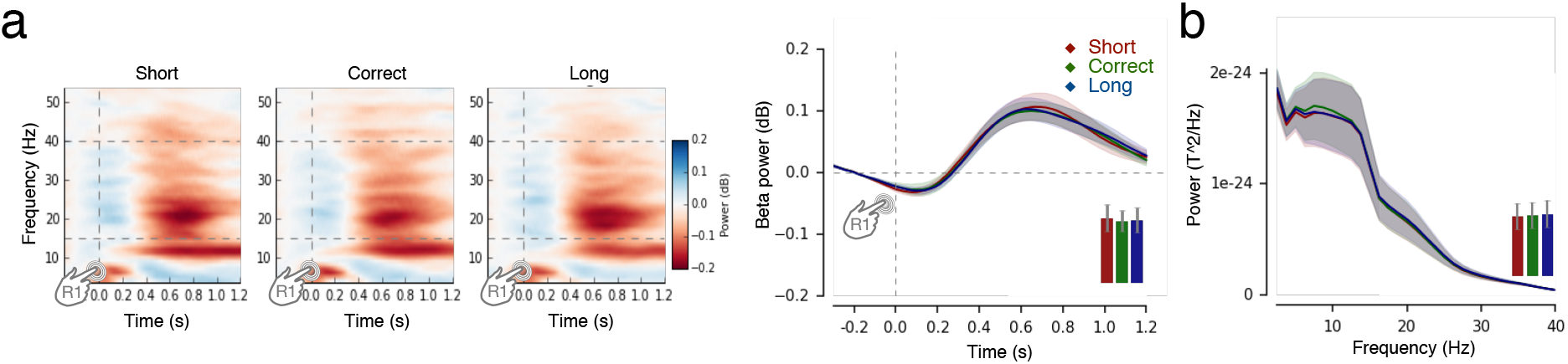
**(a)** Time-frequency spectrum for the same MEG sensor as was used for the analysis based on the FOJ split. The β band is marked by horizontal grey lines. The results of cluster-based permutation F-test performed over time did not show any differences. No significant clusters were found (all *p* > 0.1). The barplot displays SOJ driven data using the same parameters as in FOJ driven condition. (**b**) The same SOJ based analysis for PSD values. Again MEG results of cluster-based permutation F-test of PSD estimates as a function of SOJ showed no effects in any frequency band. No significant clusters where also present for analogical analyses based on EEG data (all *p* > 0.1)

**Figure S6.**
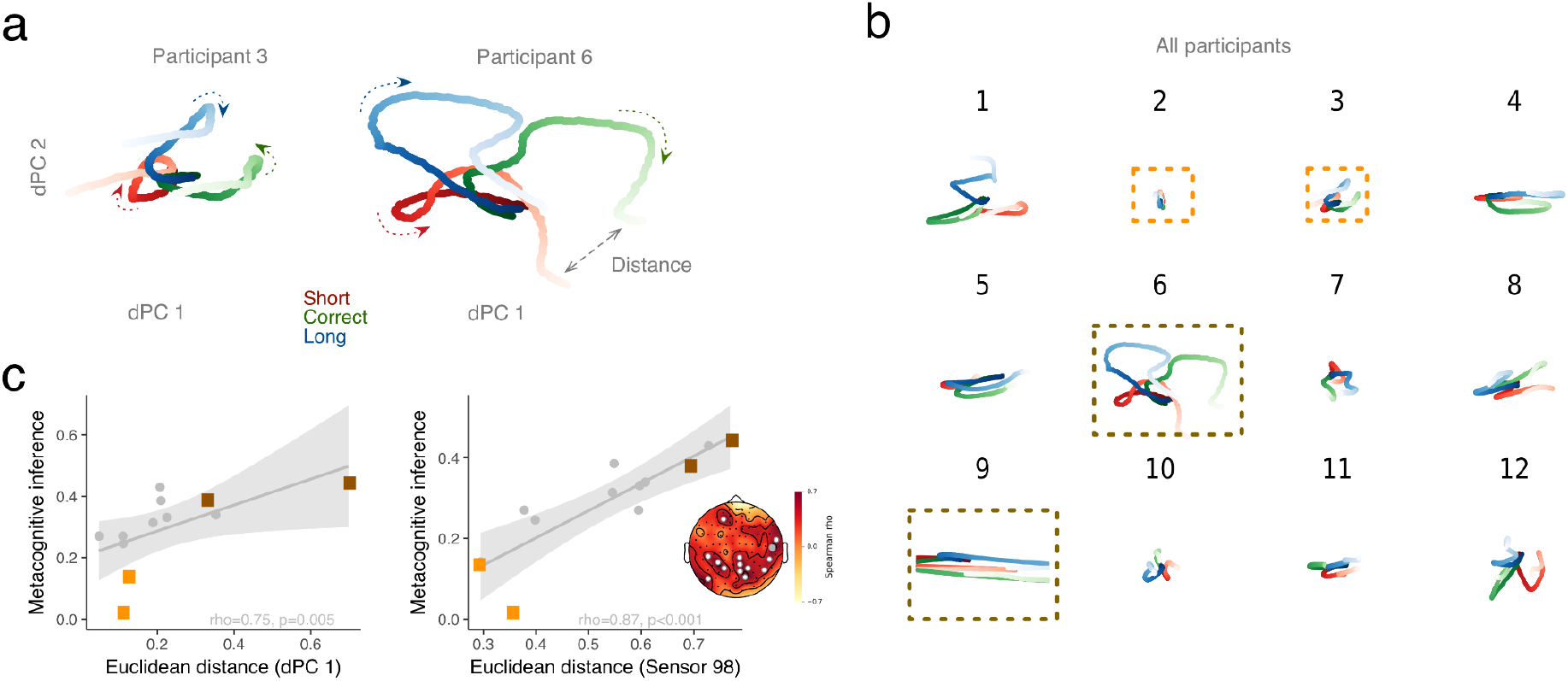
Distance in β state space supports metacognitive inference. (**a**) dPCA decomposition in β band (−0.3 to 1.2 with respect to R1) illustrated for two participants with low and high metacognitive performance, respectively. The two first dPCs are displayed against each other, and illustrate an early separation of path trajectory. Arrows indicate the progression of β state along the arrow of time. (**b)** Individual outcomes of dPCA analyses. For each participant, dPC1 was plotted against dPC2. The dark and light orange frames indicate participants with good and bad levels of metacognitive inference, respectively. (**c**) The left panel illustrates the significant correlation between individual distances in β state space (dPC1) and metacognitive inference (FOJ*SOJ) in all participants. One dot is a participant. Light and dark orange rectangles correspond to trajectories reported in (a) for two participants with a low and a high metacognitive inference ability, respectively. The right panel illustrates the significant outcome for the same analysis performed on individuals’ sensor (here, one dot is one individual, one sensor). The topographical map displays rho values together with significant sensors marked as white. The grey sensor is the one displayed in the scatterplot.

**Figure S7.**
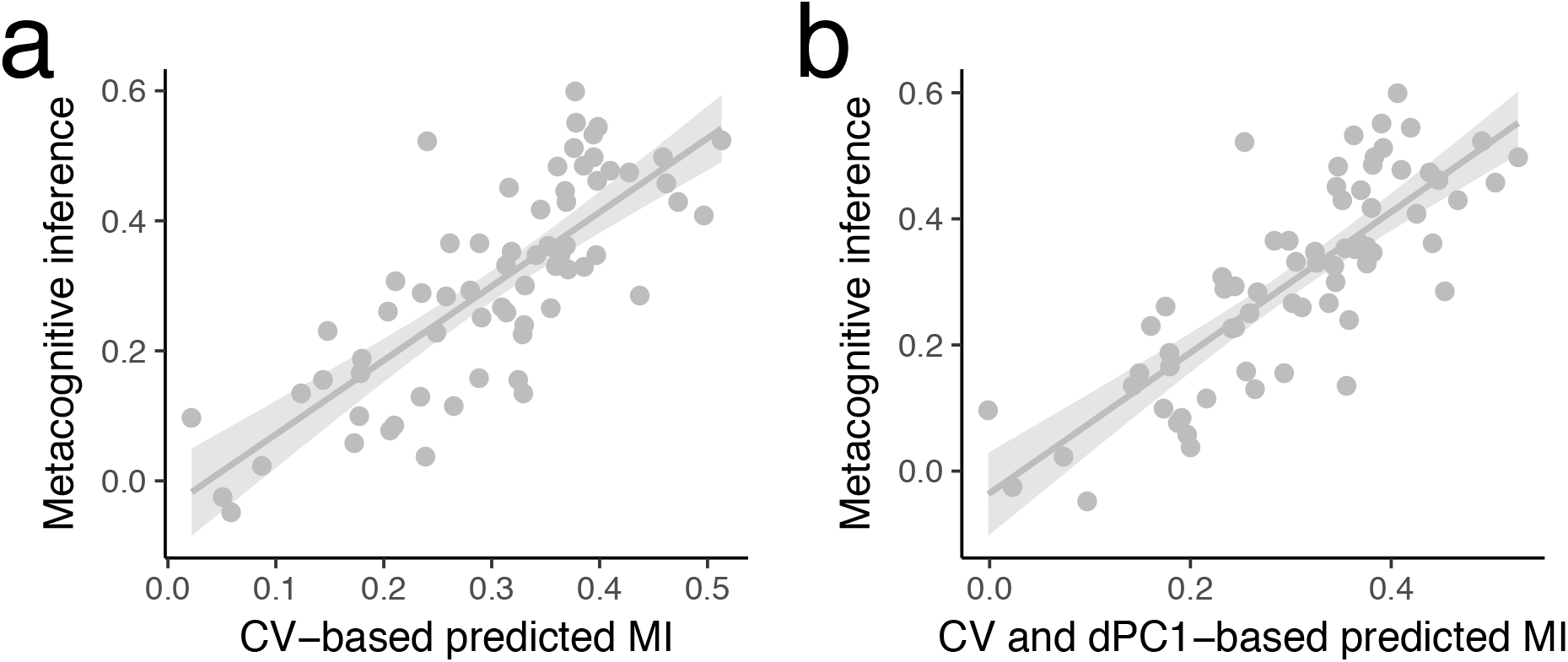
β state-space was not driven by Coefficient of Variation (CV) (Analysis per block). **(a)** A single trial analysis in which a linear mixed model was used to predict the level of metacognitive inference (FOJ*SOJ) on the basis of a Coefficient of Variation. We tested the role of behavioral fluctuations in performance precision for metacognitive inference. We assessed whether fluctuations of precision (CV) within participants contributed to the model where the distance in β state-space (dPC1) predicted metacognitive inference. **(b)** A single trial analysis in which a linear mixed model was used to predict the level of metacognitive inference (FOJ*SOJ) on the basis of a Coefficient of Variation and dPC1. We found that the addition of the precision to the model was justified [ΔAIC = 7, p = 0.003] but could not explain away all the variance as both factors of β state space distance (dPC1; t(63) = 2.5, p = 0.014) and precision (CV; t(65) = 3.1, p = 0.003) were significantly contributing to the model. The lack of correlation supported the results of our control experiment suggesting that participant did not intentionally increased their behavioral variance. Instead, the within-subject effect was indicative of the properties of internal variables. Therefore, our results bring an interesting consideration that precision on a given block may have contributed to the metacognitive performance, suggesting that participants tracked the properties of endogenous timing uncertainties (Balci et al., 2009). The parameters of internal noise could contribute to metacognitive performance, which should be closer investigated in future studies.

**Figure S8.**
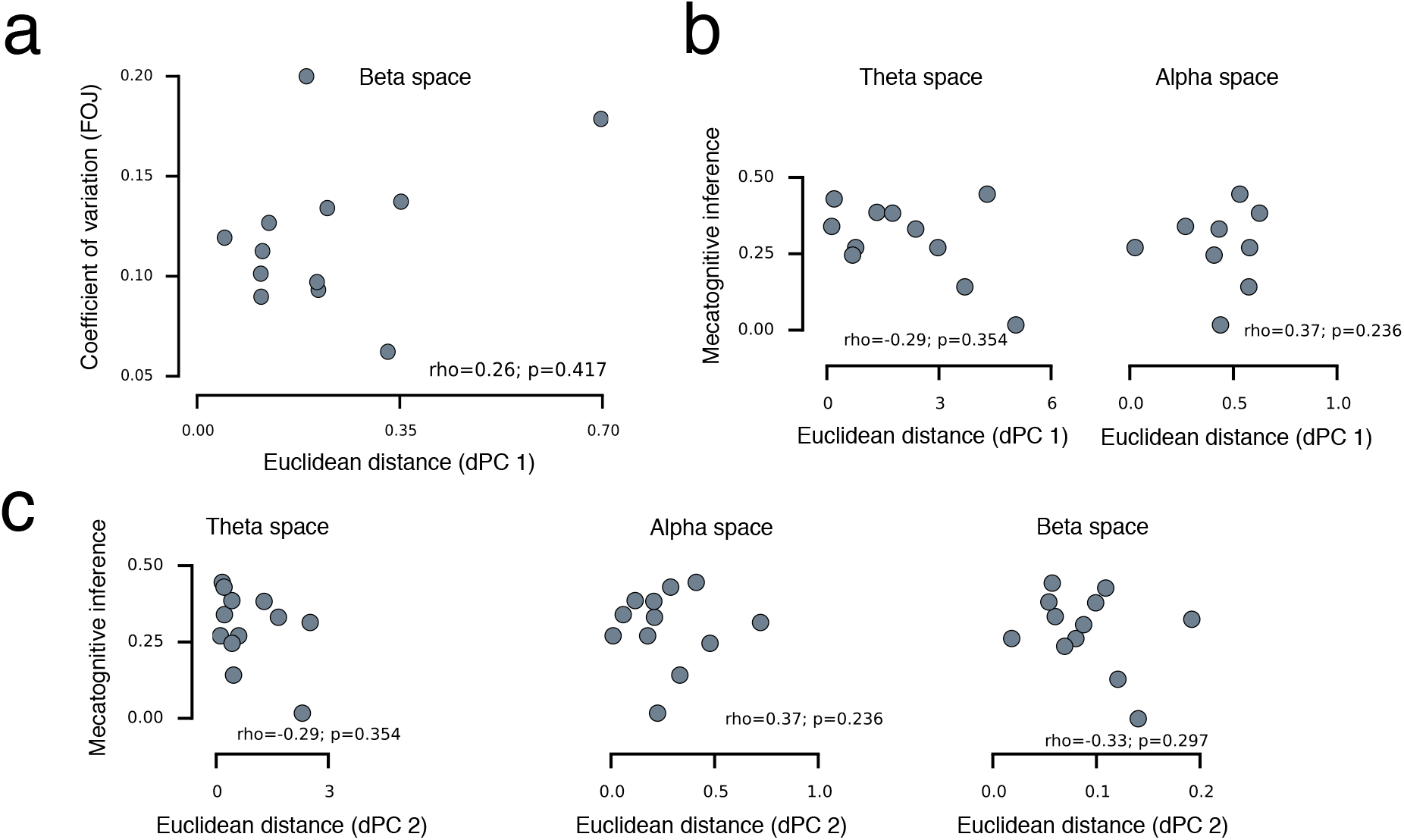
β state-space was not driven by Coefficient of Variation (CV) (Analysis per participant). **(a)** Individual distances in β state space (dPC 1) plotted against individual spread of FOJ (coefficient of variation). The multidimensional distance in β state-space was not driven by the spread of individual FOJ responses as quantified by their Coefficient of Variation (CV-FOJ; standard deviation of an individual’s FOJ divided by its mean) (*rho* = 0.26, *p* = 0.42). To insure that there was no confounds with behavioral variance in CV-FOJ, we computed robust linear regressions: a comparison of the null model to the model containing β power was justified (F_robust_ = 8.0, *p* = 0.004) whereas the inclusion of CV-FOJ against the null model was not warranted (F_robust_ = 0.03, *p* = 0.9). This demonstrated that including the CV-FOJ did not significantly account for the variance in the model. CV-FOJ and dPC1 were also dissociated as the model containing CV-FOJ and dPC1 was preferred over the model containing only CV (F_robust_ = 7.3, *p* = 0.006). The dissociation between CV-FOJ and dPC1 showed that the distance in β state-space was not solely driven by inter-individual variability in FOJ, further precluding the possibility that participants might have intentionally increase CV to obtain a better metacognitive performance. (**b**) Individual distances in theta and alpha state spaces (dPC 1) plotted against metacognitive inference (FOJ*SOJ). Although, we specifically hypothesized about β, we also insured that no other frequency bands or components were significant (all *p* > 0.1). (**c**) Individual distances in theta, alpha, and β state spaces (dPC 2) plotted against metacognitive inference.

## SUPPLEMENTAL TABLES

**Table S1.**
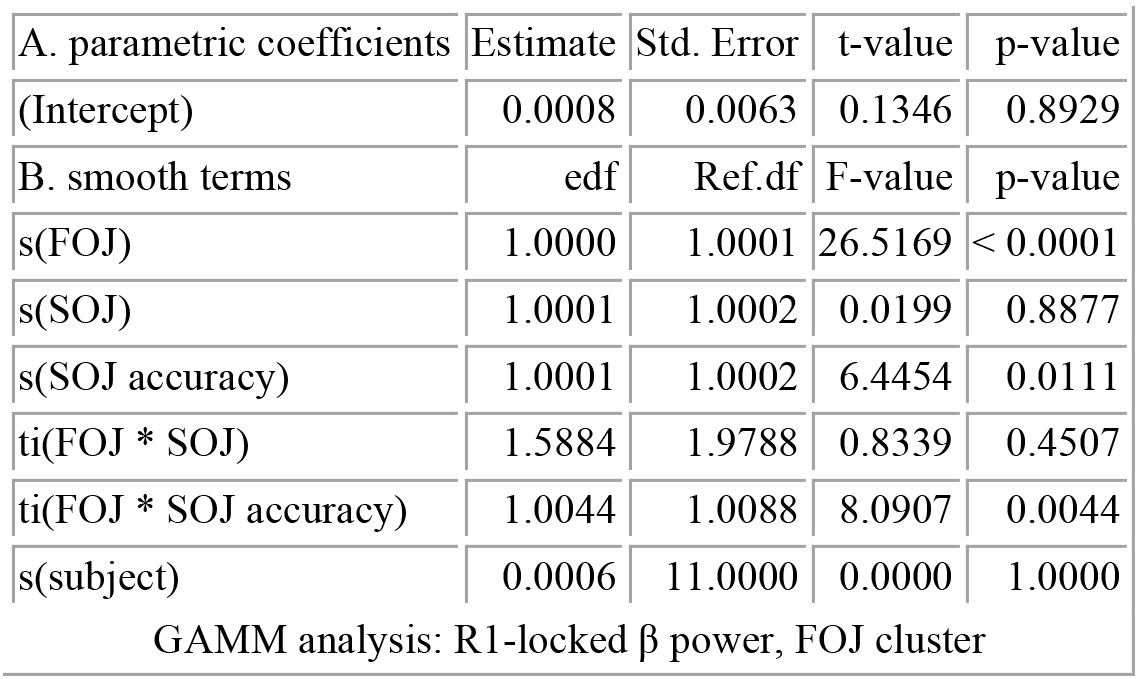
The results of single-trial GAMM analysis based on β power following the first keypress (R1). The table displays the results for the final model, which was based on the data collapsed across the significant sensors, showing the main effect of FOJ, when the model was fitted on a per sensor basis.

| A. parametric coefficients | Estimate | Std. Error | t-value | p-value |
| --- | --- | --- | --- | --- |
| (Intercept) | 0.0008 | 0.0063 | 0.1346 | 0.8929 |
| B. smooth terms | edf | Ref.df | F-value | p-value |
| s(FOJ) | 1.0000 | 1.0001 | 26.5169 | < 0.0001 |
| s(SOJ) | 1.0001 | 1.0002 | 0.0199 | 0.8877 |
| s(SOJ accuracy) | 1.0001 | 1.0002 | 6.4454 | 0.0111 |
| ti(FOJ * SOJ) | 1.5884 | 1.9788 | 0.8339 | 0.4507 |
| ti(FOJ * SOJ accuracy) | 1.0044 | 1.0088 | 8.0907 | 0.0044 |
| s(subject) | 0.0006 | 11.0000 | 0.0000 | 1.0000 |
| GAMM analysis: R1-locked $\beta$ power, FOJ cluster | | | | |

**Table S2.** The results of single-trial GAMM analysis based on β power following the first keypress (R1). The table displays the results for the final modelwhich was based on the data collapsed across the significant sensors, showing the interaction effect between FOJ and SOJ accuracy, when the model was fitted on a per sensor basis.

| A. parametric coefficients | Estimate | Std. Error | t-value | p-value |
| --- | --- | --- | --- | --- |
| (Intercept) | 0.0012 | 0.0093 | 0.1274 | 0.8986 |
| B. smooth terms | edf | Ref.df | F-value | p-value |
| s(FOJ) | 3.9387 | 5.0159 | 5.0553 | 0.0001 |
| s(SOJ) | 1.0000 | 1.0001 | 15.0954 | 0.0001 |
| s(SOJ accuracy) | 1.0001 | 1.0002 | 1.1390 | 0.2859 |
| ti(SOJ * FOJ) | 2.2156 | 2.7321 | 1.2167 | 0.4324 |
| ti(FOJ * SOJ accuracy) | 1.0018 | 1.0036 | 17.0089 | < 0.0001 |
| s(subject) | 0.0010 | 11.0000 | 0.0000 | 1.0000 |
| GAMM analysis: R1-locked $\beta$ power, FOJ * SOJ accuracy cluster | | | | |

**AUTHOR CONTRIBUTIONS**
V.W., C.R., and T.W.K. designed the research. C.R. and T.W.K. performed the experiments. T.W.K. and V.W. analyzed the data. T.W.K. and V.W. wrote the manuscript.

## ACKNOWLEDGMENTS

This work was supported by an ERC-YStG-263584, an ANR10JCJC-1904, and an ANR-16-CE37-0004-04to V.vW. We thank the members of UNIACT and the medical staff at NeuroSpin for their help in recruiting and scheduling participants. We thank members of UNICOG for fruitful discussions. Preliminary data were presented at SFN (2012, Washington DC), SFN (2015, Washington DC) and TRF (2017, Strasbourg).

